# Spatiotemporal orchestration of multicellular transcriptional programs and communications in the early stage of spinal cord injury

**DOI:** 10.1101/2022.10.07.511269

**Authors:** Zeqing Wang, Zhuxia Li, Tianle Luan, Guizhong Cui, Shunpan Shu, Yiyao Liang, Jingshu Xiao, Kai Zhang, Wei Yu, Jihong Cui, Ang Li, Guangdun Peng, Yanshan Fang

## Abstract

While spinal cord injury (SCI) involves a complex cascade of cellular and pathological changes that last for months to years, the most dramatic and comprehensive molecular rewiring and multicellular re-organization occur in the first few days, which determine the overall progression and prognosis of SCI, yet remain poorly understood. Here, we resolved the spatiotemporal architecture of multicellular gene expression in a mouse model of acute SCI, and revealed the coordinated gene co-expression networks, the upstream regulatory programs, and *in situ* cell-cell interactions that underlay the anatomic disorganization as well as the immune and inflammatory responses conferring the secondary injury. The spatial transcriptomic analysis highlights that the genes and cell types in the white matter (WM) play a more active and predominant role in the early stage of SCI. In particular, we identified a distinct population of WM-originated, *Igfbp2*-expressing reactive astrocytes, which migrated to the grey matter and expressed multiple axon/synapse-supporting molecules that may foster neuron survival and spinal cord recovery in the acute phase. Together, our dataset and analyses not only showcase the spatially-defined molecular features endowing the cell (sub)types with new biological significance but also provide a molecular atlas for disentangling the spatiotemporal organization of the mammalian SCI and advancing the injury management.

## MAIN

Spinal cord injury (SCI) causes neuronal death, synapse loss, axon retraction, demyelination, inflammation and immune activation, macrophage and immune cell infiltration, gliosis and so forth, which may result in permanent motor, sensory and autonomic dysfunctions accompanied by various local or systemic complications. Both the primary injury and the secondary injury (caused by the initial injury-induced responses) contribute to the overall pathology and disease progression^1-4^. While the chronic phase of SCI can last for months to years, the most dramatic and intensive changes in gene expression and molecular programming take place in the immediate phase (within hours) and the acute phase (a couple of days), followed by the subacute phase (the first two weeks) when secondary injury is triggered and glial scar forms, which leads to further damage to the spinal cord and sets a barrier to axonal regeneration in the chronic phase^1,5,6^. Thus, dissecting the multicellular transcriptional programs and cell-cell communications in the early stage of SCI, in particular the immediate and the acute phases that foretell the injury site re-organization and the overall prognosis, is instrumental to the understanding of SCI pathology and development of new therapeutic strategies.

Significant advancements and valuable insights into cell type-specific regulatory programs of gene expression in SCI have been achieved with the advent of single-cell (sc) and single-nucleus (sn) RNA-sequencing (RNA-seq)^7-9^. However, a spatially-resolved view of the changes in the transcriptional landscape and the molecular anatomy of the different spinal cell types as well as their context-dependent injury responses are largely missing, especially for the “enigmatic” early stage of SCI. The recent innovation in spatial transcriptomics, which allows high-throughput measurement of gene expression while retaining the anatomic information^10,11^, has been applied to build cell atlases and characterize different biological processes, including normal tissue and organ development, pathological and neurodegenerative diseases^12-21^. Given its value in molecular profiling of physically intact tissue samples in chronic conditions, we made an attempt to employ spatial transcriptomics to investigate traumatically damaged mouse spinal cords in acute injury, aiming to gain a holistic understanding of the intertangled early-phase transcriptional programs and regulatory networks that determine and underlie the SCI-induced molecular and cellular changes in their morphological context.

### The mouse SCI model and the workflow of the spatial transcriptomic analyses

For the spatial transcriptomic analysis of mammalian SCI in this study, the spinal cord of C57BL/6J mice was fully transected at the tenth thoracic vertebra (T10) (Fig. 1a and Extended Data Fig. 1a). The motor function and the morphology of the spinal cord after injury were examined at 0 (uninjured), 3, 24 and 72 hours post injury (hpi) to confirm a complete transection (Extended Data Fig. 1b-d). We performed hematoxylin and eosin (H&E) staining on transverse sections of various distances from the injury epicenter on both the rostral (-) and caudal (+) segments (Extended Data Fig. 1e). As shown in Extended Data Fig. 1d and e, the spinal cords were severely deformed within ±0.25 mm, precluding an informative spatial transcriptomic analysis. Therefore, the rostral (-) and the caudal (+) sections at ±0.5 and ±1.0 mm from the injury epicenter were subjected to the 10x Visium spatial RNA-seq analysis. Four consecutive cryosections each at ±0.5 and ±1.0 mm were captured, and the spinal sections of the same distance at different time points (0, 3, 24 and 72 hpi) were attached to the same Visium Gene Expression slide and processed simultaneously in the subsequent procedure (Fig. 1b). Together, 64 spinal cord sections from 9 different mice were examined in this spatial transcriptomic analysis, representing an inclusive coverage of the dynamic injury responses in a high spatial and temporal resolution.

**Fig. 1.**
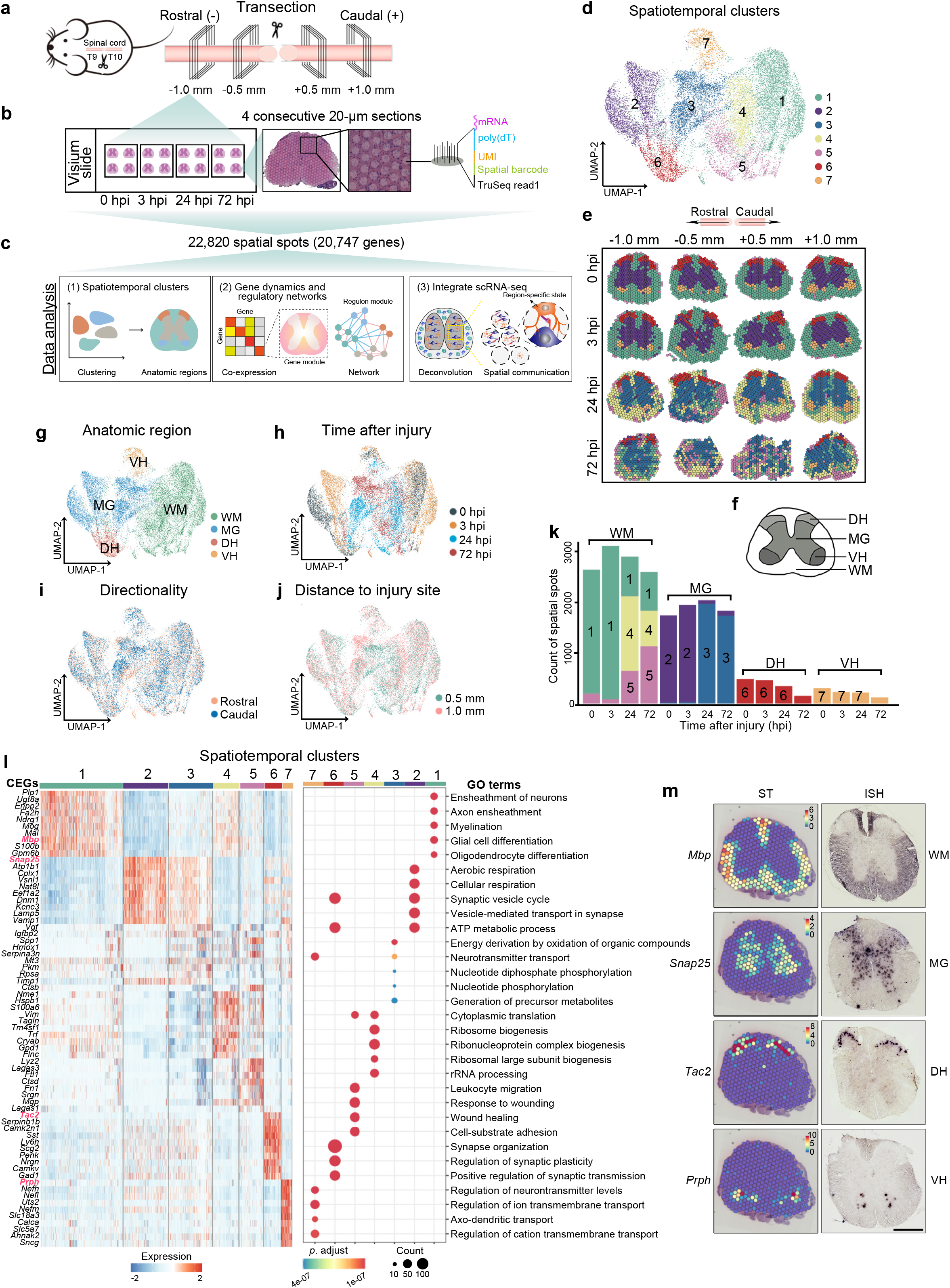
Spatiotemporal transcriptomics and molecular clusters of mouse SCI using Visium. (**a**) A carton of the mouse SCI model and the spinal cord sections obtained in this spatial transcriptomic study. (**b**) A schematic of the Visium Gene Expression slide and the acquisition of the mouse SCI sections for RNA-sequencing. (**c**) The main workflow of the spatial transcriptomic data analyses. (**d**) The UMAP plot of the 22,820 transcriptomic profiles (one profile for each spatial spot) of all the SCI samples identified 7 spatiotemporal clusters. (**e**) The clusters are color coded and shown in the spatial view of the spinal cord sections at the indicated locations and time points after injury. (**f**) A cartoon showing the main anatomic regions of the spinal cord: WM, white matter; MG, middle grey; DH, dorsal horn; VH, ventral horn. (**g-j**) The UMAP plots of the spatial spots were colored according to anatomic region (**g**), time after injury (**h**), directionality (**i**) and distance to the injury site (**j**). (**k**) Quantification of the numbers of the spatial spots in each cluster at the indicated time points. (**l**) The heatmap showing the expression levels of the top 10 CEGs (left) and the dot-plot showing the top 5 GO terms (right) for each of the 7 spatiotemporal clusters. (**m**) The spatial view of the gene expression levels in the spatiotemporal transcriptomics (ST, left) and *in situ* hybridization (ISH, right) of four representative anatomic markers as indicated. Scale bar: 500 μm. SCI, spinal cord injury; hpi: hours post injury; UMI, unique molecular identifier; UMAP, uniform manifold approximation and projection; CEGs, cluster-enriched genes; GO terms, gene ontology terms; *p*. adjust, adjusted *p* value.

Overall, 22,820 spatial spots were obtained and profiled after filtering, with a mean of 12,570 RNA reads and 3,996 genes per spot (Extended Data Fig. 2a-d and Supplemental Table 1). The discernible grey matter (GM) regions containing condensed neuronal cells showed higher numbers of expressed genes (Extended Data Fig. 2d). We compared the intra- and inter-correlations of the sample groups and found a high data reproducibility of the replicate sections within each group at the bulk level (Extended Data Fig. 2e). We therefore analyzed all the sampled spots of the four replicates but visualized the results on each representative sample section in all the spatial view charts in this study.

Further spatiotemporal evaluation of the expression of known injury-related genes (e.g., *Atf3, Hmox1* and *Timp1*), aka, the injury score (see Methods), displayed remarkable patterns in association with the injury time, the distance to injury epicenter and the anatomy of the spinal cord (Extended Data Fig. 2f, g). For example, the ±0.5 mm spinal sections at 72 hpi displayed the highest injury scores (Extended Data Fig. 2f, g) and the lowest transcriptomic similarity to others (Extended Data Fig. 2e), which were in accordance with the most severe damage among all the 64 sections (Extended Data Fig. 1e). Therefore, in the subsequent data analyses, we focused on revealing: (1) the spatiotemporal molecular annotation of the spinal cord anatomy, (2) the gene co-expression modules and regulatory networks, and (3) the spinal cell type composition and cell-cell communications in response to SCI (Fig. 1c). For readers with broader research interests, an interactive data exploration portal is available at: *(will release to the public upon acceptance of the paper)*.

### Cluster analysis of the spatial spots reveals anatomic domain- and injury time-featured expression profiles of SCI

We first performed dimension reduction and cluster visualization on the collective spatial spots using uniform manifold approximation and projection (UMAP) (see Methods). Seven distinct gene expression clusters were identified (Fig. 1d) and Clusters 1, 2, 6 and 7 showed a bona fide anatomic and molecular annotation of the intact spinal cord at 0 and 3 hpi (Fig. 1e). This allowed us to categorize the seven clusters into four spatial domains according to the four major anatomic regions of the spinal cord: white matter (WM), middle grey (MG), dorsal horn (DH) and ventral horn (VH) (Fig. 1f, g). With the injury score increased along the time (Extended Data Fig. 2f, g), the physical boundary of the anatomic regions became less clear at 72 hpi especially with the ±0.5 mm sections (Fig. 1e). However, the cluster analysis could still classify the spatial spots to the four distinct spatial domains following the coherence in the UMAP plot (Fig. 1g), which showcased the advantage of spatial transcriptomics in providing the molecular annotation of tissue architecture to better understand the SCI pathology.

Time after injury, along with the anatomic domain, comprised the major transcriptomic differences amongst the spatial spots of the spinal cord examined in this study (Fig. 1h). In contrast, the rostral/caudal directionality and the distance to the injury site showed mild impacts on distinguishing the clusters (Fig. 1i, j). Consistently, analysis of differentially expressed genes (DEGs) revealed remarkable domain-dependent alterations, whereas only a small number of directionality-or distance-specific DEGs were identified during SCI (except for the ±0.5 mm sections at 72 hpi, where more severe damage was associated) (Extended Data Fig. 3a-c). As such, we focused on the factors of spatial domain and injury time in the subsequent analyses (also see Extended Data Fig. 3d-g).

A remarkable shift of the compositions of spatiotemporal clusters in the WM and MG domains was elicited by SCI (Fig. 1k). To understand the molecular changes and functional relevance of these changes, we identified the cluster-enriched genes (CEGs) for each cluster and analyzed their spatiotemporal expression profiles (Fig. 1l). The CEGs included not only the well-known markers of the four anatomic regions of the intact spinal cord, such as *Mbp* (Cluster 1) for WM, *Snap25* (Cluster 2) for MG, *Tac2* (Cluster 6) for DH and *Prph* (Cluster 7) for VH, with their spatial expression validated by *in situ* hybridization (ISH) (Fig. 1m), but also uncharted signature genes for the different spatial domains especially after injury (Extended Data Fig. 4). Gene ontology (GO) term enrichment analyses of the CEGs revealed the biological processes that not only reflect the unique physiological function of the different anatomic regions, but also underlie the pathological progression of SCI. For example, Cluster 1 was predominantly localized to the WM at 0 and 3 hpi (Fig. 1k), and its CEGs, such as the myelin-related genes *Plp1, Mog* and *Mbp*^22^, were enriched in the biological functions of myelination and oligodendrocyte differentiation (Fig. 1l). At 24 and 72 hpi, the main clusters of the WM switched to Clusters 4 and 5 (Fig. 1k), whose CEGs, such as the reactive astrocyte markers *Vim* and *Tm4sf1*^23^ in Cluster 4 and the activated macrophage markers *Lyz2* and *Lgals3*^24^ in Cluster 5, were enriched with the terms of protein synthesis, cell migration, responses to wounding, *etc*. (Fig. 1l). Thus, the switch of the spatiotemporal clusters in the WM between 3 and 24 hpi suggested a reduction of myelination-related functions along with the activation of astrocytes and infiltration of myeloid cells in the WM regions between the immediate phase and the acute phase of SCI.

### Spatiotemporal analysis of gene co-expression patterns and transcriptional programs in SCI

Next, to depict the intricate and dynamic transcriptional changes in SCI and unveil the commonalities in the molecular events and regulations, we performed the weighted gene co-expression network analysis (WGCNA)^25^ (Fig. 2a). Thirty-three spatially and temporally co-expressed gene modules (CGMs) were identified (see Methods and Supplemental Table 2), which displayed unique and diverse spatiotemporal expression patterns, cellular functions, gene connectivity networks and transcriptional regulations (Extended Data Fig. 5 and 6), together providing an unbiased molecular annotation of the injury-induced expression programs in SCI. For example, CGM13 showed an injury-induced expression in the WM specifically at 3 hpi, after which the expression was sharply decreased (Fig. 2b). This module was highly associated with the cellular functions of regulating the ERK1/2 signaling cascade, and the node genes of the connectivity network were *Csrnp1* and *Maff*, which were known players in stress, immune and inflammatory responses^26,27^. Thus, CGM13 likely represented a group of MAPK/ERK-mediated “early response genes” regulating immunity and inflammation in the acute phase of SCI.

**Fig. 2.**
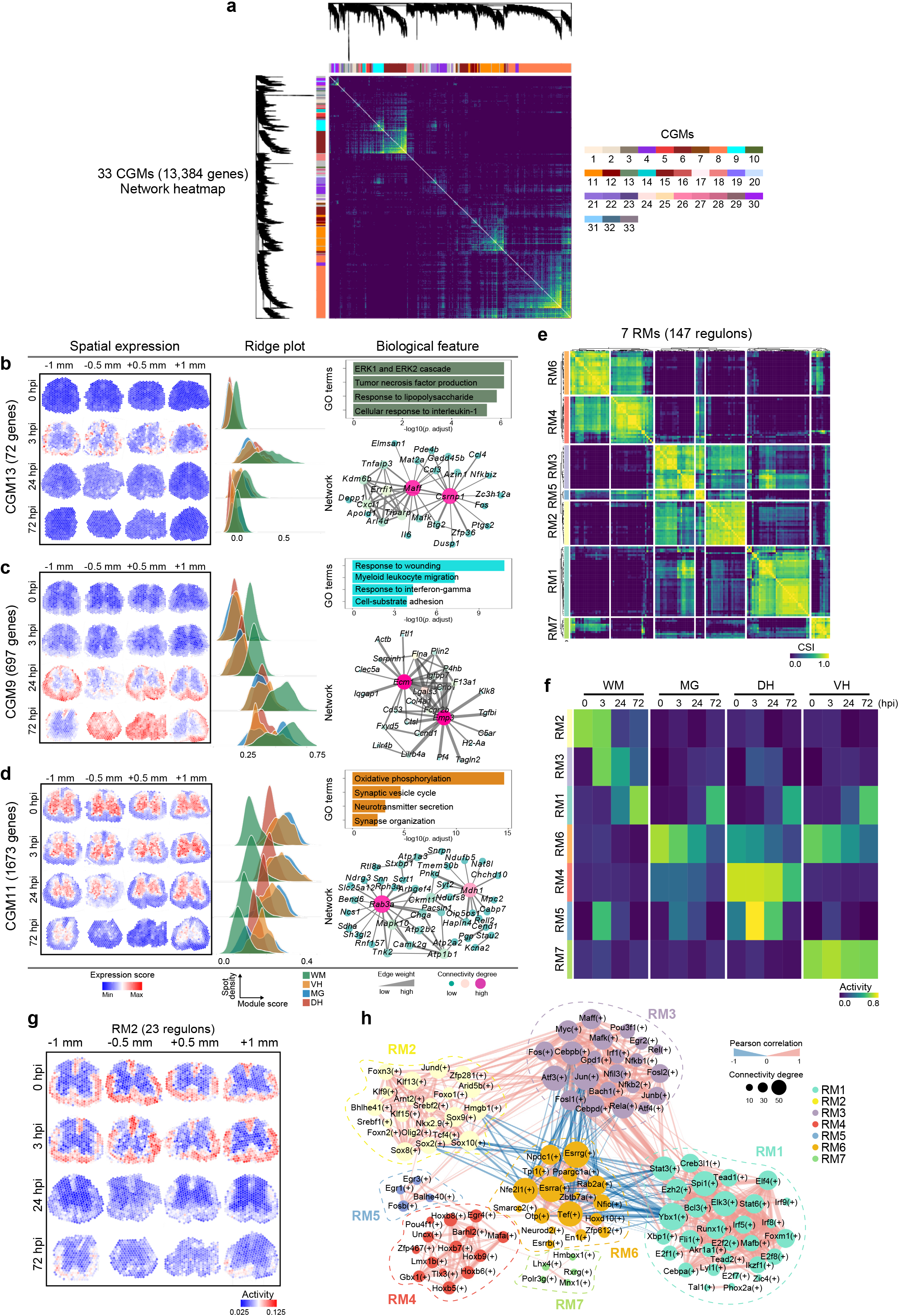
Spatiotemporal analyses of gene co-expression dynamics and regulatory networks in SCI. (**a**) Heatmap showing the weighted-correlation network of the genes constituting 33 SCI spatiotemporal CGMs. (**b-d**) The molecular features of three representative CGMs: CGM13 (**b**), CGM9 (**c**) and CGM11 (**d**). From left to right: Left, the spatial view of average levels of the genes expressed in the indicated CGM during the process of SCI; middle, the ridge plot quantitatively illustrating the density distribution of the CGM expression within the 4 anatomic domains at the indicated time points of SCI; right, the representative GO terms (top) and the gene network showing the highest 50 connectivity weights (bottom) of the indicated CGM. The number of genes in each CGM is shown in the parentheses. (**e**) The CSI heatmap of 147 transcriptional regulons grouped into 7 RMs by hierarchical clustering. (**f**) The heatmap of the overall activity of each RM enriched in the four anatomic domains at the indicated time points of SCI. (**g**) The spatial view of the average regulon activity of RM2 at the indicated time points of SCI. The number of regulons in RM2 is shown in the parentheses. (**h**) The Pearson correlation networks and the CSI connectivity degree of the transcriptional regulons within and across the 7 RMs. CGM, co-expressed gene module; CSI, connectivity specificity index; RM, regulon module.

In addition, we noticed that some modules, such as CGM9 and CGM11, exhibited an interesting spatiotemporally-complementary expression pattern: CGM9 was expressed at a low level in the WM of the intact spinal cord (0 hpi) and at 3 hpi, but was robustly increased at 24 hpi, which expanded into the MG in the ±0.5 mm sections at 72 hpi (Fig. 2c). In contrast, CGM11 was highly expressed in the MG at 0 and 3 hpi, which started to decrease at 24 hpi and only retained weak expression at 72 hpi, with the expression in the ±0.5 mm sections reduced the most (Fig. 2d). Scrutinizing the biological functions and the connectivity network of the genes enriched in these two CGMs, we found that CGM9 was mainly involved in stress, wound and immune responses, whereas CGM11 was mostly associated with the function and maintenance of neuronal synapse. Thus, this complementary expression pattern of the two CGMs likely underscored the interplay of SCI-induced immunity and inflammation with the loss of neurons and synapses during SCI.

To further unravel the regulatory mechanisms that determine the spatiotemporal changes in gene expression in SCI, we performed single-cell regulatory network inference and clustering (SCENIC) analysis^28^ to identify transcriptional regulons (consisting of the transcription factors (TFs) and their target genes) that could coordinately regulate gene expression in response to SCI. Overall, 147 regulons were identified and classified into seven regulon modules (RMs) (Fig. 2e and Supplemental Table 3). The RMs showed remarkable anatomic domain- and/or injury time-specificity (Fig. 2f and Extended Data Fig. 7). For example, RM4 and RM7 functioned predominantly in the DH and the VH, respectively; RM6 was enriched in the GM regions including MG, DH and VH and its activity was gradually reduced along the time in SCI; RM5 was highly expressed in the DH and the WM at 3 hpi, which may be associated with the induction of the early response genes.

The RMs of the WM were more complicated and dynamic: RM2 was the main RM at 0 and 3 hpi, and it was superseded by RM3 at 3 and 24 hpi and finally by RM1 at 72 hpi (Fig. 2f, g and Extended Data Fig. 7a, b). Along with these changes, the TFs of the main regulons in the WM switched from Sox10, Oligo2 and Srebf2 (RM2), to Fos, Jun and Nfkb1/2 (RM3), and ultimately to Stat3, Stat6 and Irf5 (RM1) (Fig. 2h). Sox10 and Oligo2 are important TFs in oligodendrocyte differentiation and maturation^29^ and Srebf2 regulates genes involved in cholesterol biosynthesis^30^; Fos and Jun are key TFs in TGFβ-mediated signaling and regulate cell proliferation, differentiation and death^31^ and Nfkb1/2 is a central activator of genes related to inflammation and immune functions^32^; Stat3, Stat6 and Irf5 mediate gene expression in response to cell stimuli and are widely involved in immunity, inflammation and cell death^33^. Together, the shifts of the main RMs in the WM (Fig. 2f) were in agreement with the changes of the spatiotemporal clusters in the WM during SCI (Fig. 1k), which underscored the injury-induced oligodendrocyte dysfunction and immune and inflammation activation in the WM. Of note, the immune and inflammatory responses mediated by RM3 at 3 and 24 hpi were largely restricted to the WM, whereas those by RM1 at 72 hpi were comparably activated in all the four anatomic domains (Fig. 2f and Extended Data Fig. 7a), suggesting a widespread upregulation of immune and inflammatory genes by the end of the acute phase of SCI.

### Integration of the SCI scRNA-seq data for an *in situ* multicellular gene expression and cell communication atlas

Because the Visium spot is not single-cell resolution, to allocate the cell type information to the SCI spatial spots in a comparable manner, we performed deconvolution analysis by robust cell type decomposition (RCTD^34^; also see Methods) using the combined normal and injured spinal cord scRNA-seq datasets as reference^9,35^. The proportions of different cell types detected within a single spatial spot were displayed as a pie chart and the entire spinal cord section was displayed as an assembly of the spatial pie charts (Fig. 3a, b). The overall composition of the main cell types and their anatomic localizations were in good accordance with the spatial architecture of the spinal cord. For example, neurons and oligodendrocytes were predominantly localized to the GM and the WM, respectively (Fig. 3b). Interestingly, we found that the two molecularly-defined astrocyte subtypes, astrocyte-*Gfap* and astrocyte-*Slc7a10*^35^ (Supplemental Table 4), showed distinct and mutually exclusive distributions in the white (namely, astrocyte-WM) and grey matter (namely, astrocyte-GM), respectively (Fig. 3b), which represented the general categories of astrocyte heterogeneity in location of white (fibrous astrocyte) versus grey matter (protoplasmic astrocyte)^36^. Furthermore, deciphering *in situ* spinal cell type compositions and their spatiotemporal dynamics during SCI revealed sophisticated spatial domain- and injury time-associated alterations (Extended Data Fig. 8), including the emergence of the cell types that participate in the scar formation such as fibroblasts, ependymal and vascular cells as well as the invasion of peripheral immune and inflammatory cell types such as myeloid cells, macrophages, monocytes and neutrophils (Fig. 3c and Extended Data Fig. 9).

**Fig. 3.**
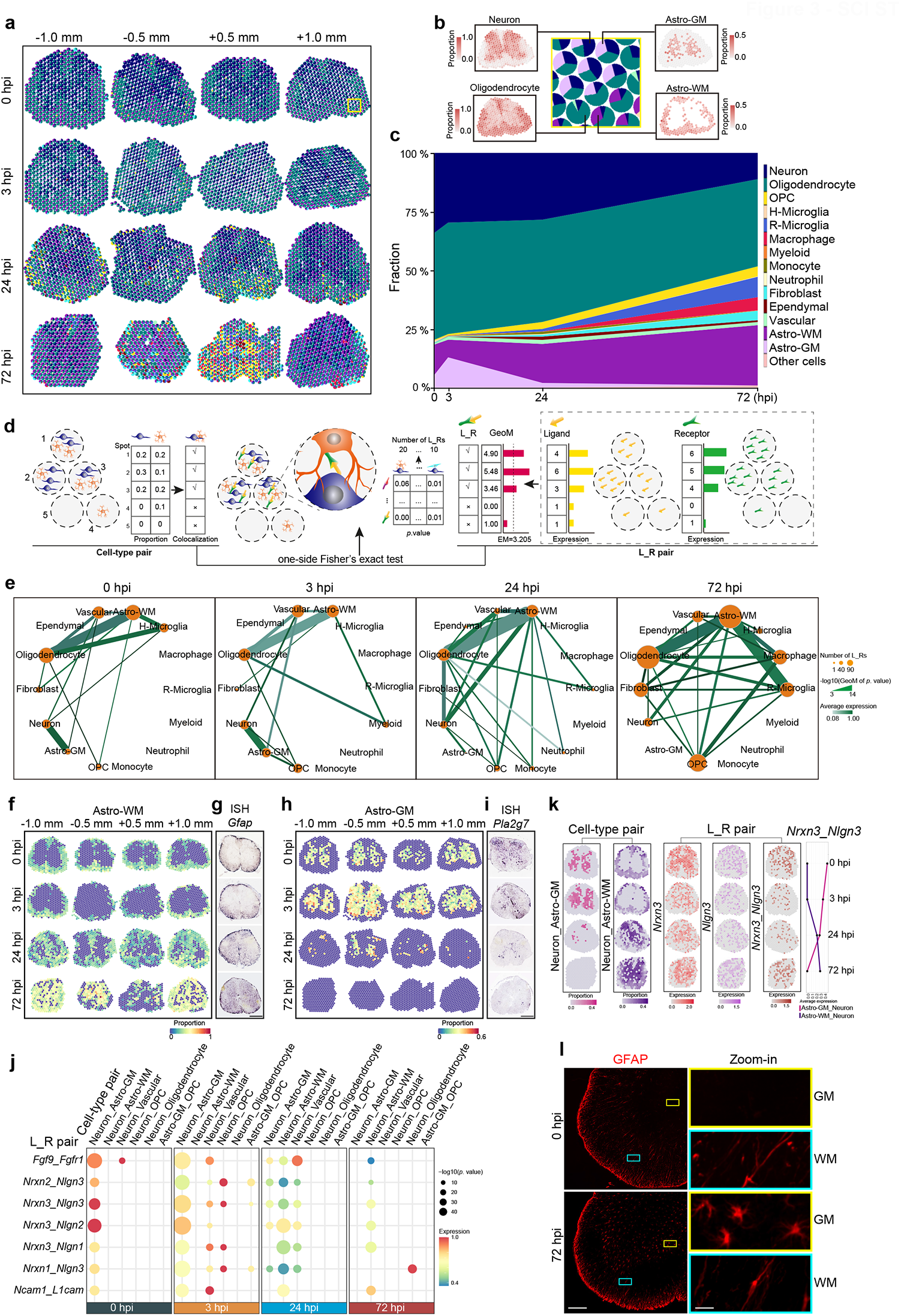
Analyses of multicellular responses and cell-cell communications in SCI identify injury-induced, GM-relocated astrocytes-WM. (**a, b**) The spatial views of the spots showing the proportions of all cell types in each spot (as a pie chart) of the spinal sections at the indicated time points in SCI (**a**), and the zoom-in of the yellow-boxed area with the spatial distributions of the representative spinal cord cell types is shown in (**b**); also see Extended Data Fig. 7-8. (**c**) The fractions (%) of different cell types at indicated time points in SCI. (**d**) A schematic cartoon illustrating the main procedure for the spatial-aware cell-cell communication (SA-CCC) analysis that evaluates *in situ* intercellular communications by testing the statistical significance of the co-occurrence of a cell-type pair and an L_R pair. (**e**) The SA-CCC networks at each indicated time point in SCI. The size of the orange circles represents the number of the interacting L_R pairs expressed in each cell-type pair. For the connecting lines, the thickness denotes the GeoM of the significance of the interaction and the shade indicates the average expression levels of the L_R pairs for each cell-type pair. (**f-i**) The spatiotemporal distributions of astrocytes-WM (**f**) and astrocytes-GM (**h**), validated with ISH of *Gfap* for astrocytes-WM (**g**) and *Pla2g7* for astrocytes-GM (**i**). Scale bars: 500 μm. (**j**) Dot plots showing the major interacting L_R pairs of the main cell-type pairs in the GM at the indicated time points in SCI (from Extended Data Fig. 10a). (**k**) From left to right: left, the geometric proportions of the cell-type pairs of “astrocyte-GM_neuron” and “astrocyte-WM_neuron”; middle, the expression levels of the individual ligand, receptor, and their geometric mean of the L_R pair of *Nrxn3*_*Nlgn3*; right, the average expression of the *Nrxn3*_*Nlgn3* pair within the indicated co-localized cell-type pairs along the time in SCI. (**l**) Immunostaining of GFAP in the mouse spinal cord (-1.0 mm section) at 0 and 72 hpi. The morphologies of GFAP^+^-astrocytes in the GM (yellow box) and the WM (cyan box) are shown in the zoom-ins on the right. Scale bars: 200 μm (left) and 20 μm (right). Astro, astrocytes; WM, white matter; GM, grey matter; H-Microglia, homeostatic microglia; R-Microglia, reactive microglia; GeoM, geometric mean; EM, assembly mean; L_R pair, ligand_receptor pair; ISH, *in situ* hybridization.

Cell-cell interaction plays important roles in maintaining tissue integrity and normal functions. To denote faithful *in situ* cell-cell interactions from our spatial transcriptomic data, we performed a spatial-aware cell-cell communication (SA-CCC) analysis. In brief, spatial colocalization of cell-type pairs within spots and spot-enriched ligand_receptor (L_R) pairs deduced from CellChat package^37^ were subjected to co-occurrence test (Fig. 3d and see Methods). Upon injury, both the number of interacting L_R pairs per cell-type pair and the complexity of the network of co-localized cell types were markedly increased, especially at 72 hpi (Fig. 3e). Among all the cell types, oligodendrocytes and astrocytes had the most extensive connections and served as the nodes to interact with other cell types in both intact and injured spinal cords, especially at 72 hpi. In addition, the connectivity of macrophages, reactive microglia, OPCs and fibroblasts was substantially increased, indicating much escalated diversity of cell-cell communications in this stage of SCI.

**Fig. 4.**
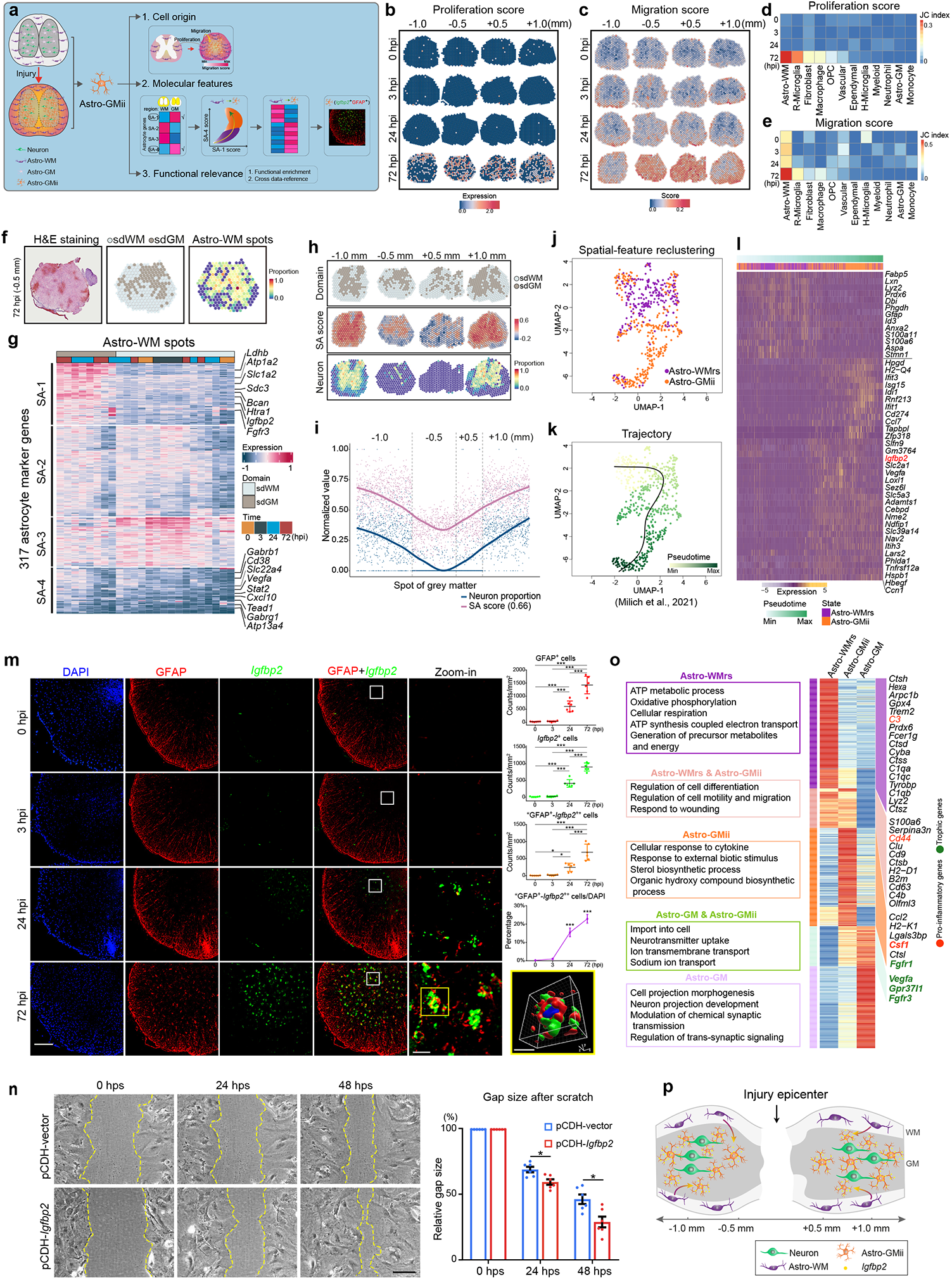
Characterization of the molecular features of SCI-induced astrocytes-GMii with spatial transcriptomics and scRNA-seq integration. (**a**) An overview of the comprehensive spatial and single-cell transcriptomic analyses (in Fig. 4 and Extended Data Fig. 12) for the identification and characterization of the SCI-induced astrocytes-GMii. (**b, c**) The spatiotemporal dynamics of the proliferation (**b**) and migration (**c**) scores in SCI. (**d, e**) The jaccard similarity (JC index) of the proliferation score (**d**) or migration score (**e**) for the spatial spots containing the specified individual cell types at the indicated time points in SCI. (**f**) An example of the spinal cord section showing the H&E staining, the spatially-defined (sd) WM and sdGM domains (annotated using the spatial clusters in Fig. 1g, k), and the spatial spots containing astrocytes-WM. (**g**) The 317 astrocyte deconvolution marker genes in the spatial spots containing astrocytes-WM are clustered into four SA groups based on their bulk expression levels according to the sdWM or sdGM domain and the injury time. (**h, i**) The spatial distributions of the sdWM and sdGM domains, the SA score and neuron proportion in each spot at 72 hpi (**h**) and their scatterplot along the distance (-1.0 to +1.0 mm) as well as the corresponding locally weighted regression fitting curves (**i**) are shown. The normalized values are the z-score [0, 1] of the SA scores and neuron proportions. The Spearman rank correlation coefficient of the SA score and neuron proportion is shown in the paratheses. (**j, k**) The UMAP re-clustering (**j**) and pseudotime trajectory (**k**) analyses of the astrocytes-*Gfap* in the SCI single-cell dataset^9^ identify distinct populations of astrocytes-WMrs and astrocytes-GMii sharing a continuous cell lineage. (**l**) The expression heatmap of the top 50 most dramatically-altered genes along the trajectory in (k). Among them, an example of *Igfbp2* is examined by FISH in (m). (**m**) Representative confocal images of immunostaining of GFAP combined with FISH of *Igfbp2* of the mouse spinal cord (-1.0 mm section) at the indicated time points in SCI. The white-boxed areas are shown in a higher magnification in the zoom-ins and the 3D rendering of a “GFAP^+^-*Igfbp2*^+^” cell at 72 hpi (yellow box) is shown, with DAPI staining to indicate the location of the nucleus. The average densities of the indicated immunophenotypes of cells in the mouse spinal cord area and the percentage of “GFAP^+^-*Igfbp2*^+^” cells to total cell counts (indicated by DAPI) at the specified time points in SCI are quantified. Scale bars: 10 μm in 3D rendering, 25 μm in zoom-ins, and 200 μm in the rest. (**n**) Representative brightfield images of the scratch assay of *in vitro* cultured mouse primary astrocytes infected with pCDH-vector (control) or pCDH-*Igfbp2*. The gap (cell-free area) caused by the scratch in multiple random visions of two pooled repeats were traced along time and the average size of the gap was calculated and shown on the right. Scale bar, 100 μm. Mean ± SEM; n = 6; \**p* < 0.05 and \*\*\**p* < 0.001; one-way ANOVA in (m) and Student’s t-test in (n). (**o**) GO term analysis of the unique and shared DEGs (left) and the heatmap showing the average normalized expression levels of the DEGs (right) associated with astrocytes-WMrs, -GMii and -GM. The DEGs that are cross-referenced as pro-inflammatory (red), neurotrophic (green) or neuroprotective (in bold) genes by Kaufmann et al.^46^ are highlighted. (**p**) A schematic of the SCI-induced, *Igfbp2*-expressing astrocytes-GMii, which originate from reactive astrocytes in the WM and migrate into the GM to provide support to spare neurons. Astro, astrocytes; WM, white matter; GM, grey matter; Astro-GMii, injury-induced GM-relocated astrocytes; Astro-WMrs, the rest astrocytes-WM; SA, spatiotemporally-distinct astrocyte; FISH, fluorescence ISH; hps, hour post scratch.

To pinpoint the specific L_R pairs that mediate the spatiotemporal dynamics of cell-cell communications, the top L_R pairs in each cell-type pair at different time points in SCI were identified and examined (Extended Data Fig. 10a) and three main classes of L_R pairs were discerned (Extended Data Fig. 10b). Unlike the cell-cell communication analysis based solely on scRNA-seq datasets, the SA-CCC analysis is featured by identifying both the expression levels and the *in situ* interacting cell types for a particular L_R pair in a context-dependent manner. For example, the L_R pair of *Spp1*_*Cd44* was expressed predominantly in the WM at moderate levels at 0 and 3 hpi, which was increased remarkably at 24 hpi (primarily in the WM and scattered in some GM regions) and spread throughout the entire spinal cord section at 72 hpi (Extended Data Fig. 10c). Accompanying this profound induction, the interactions of the cell-type pair(s) mediated by the L_R pair of *Spp1*_*Cd44* expanded from “oligodendrocyte_astrocyte-WM” to “astrocyte-WM_OPC” in the WM at 24 hpi and finally to “oligodendrocyte_macrophage” and “OPC_macrophage” in the GM at 72 hpi (Extended Data Fig. 10c), suggesting a multifaceted role of this L_R pair along the progression of SCI.

### Injury induces redistribution and repurposing of astrocytes-WM into the GM of the spinal cord

Astrocytes are morphologically and functionally diverse and play vital roles in the nervous system in both physiological and pathological conditions^38^. Being a significant portion in the spinal cord, not only the proportions of the two main astrocyte subtypes (WM and GM) altered dramatically (Fig. 3c), but also their weights in coordinating cell-cell communications in SCI differed a lot (Fig. 3e). Indeed, the gene expression features, GO term analysis of the DEGs and their upstream transcriptional regulons as well as cross-reference with two single-cell SCI datasets all pointed to the molecular basis underlying their spatial and functional divergences (Extended Data Fig. 11). Thus, our spatial classification of astrocytes-WM and astrocytes-GM corroborates the different physiological functions of the known astrocyte subtypes in the mammalian nervous system^39,40^.

Strikingly, the two astrocyte subtypes displayed quite different injury-induced changes in their spatiotemporal distributions in response to SCI: astrocytes-WM were largely unchanged at 3 hpi but were drastically increased at 24 and 72 hpi, at which they were even present in the GM region; astrocytes-GM were increased at 3 hpi but then drastically decreased at 24 hpi and became almost undetectable at 72 hpi (Fig. 3f, h). The dynamic changes of distribution were validated by examining the spatiotemporal expression patterns of two representative marker genes (*Gfap* and *Pla2g7*; see Extended Data Fig. 11e) by ISH (Fig. 3g, i). Furthermore, by the SA-CCC analysis of cell-cell interactions in the GM and the L_R pairs mediating these interactions (e.g., the *Nrxn*_*Nlgn* families) (Fig. 3j), we revealed a clear switch of neuron-interacting cells from “astrocytes-GM” at 0 and 3 hpi to “astrocytes-WM” at 24 and 72 hpi, accompanied by the synchronized switch of the spatial L_R pairs (e.g., *Nrxn3*_*Nlgn3*) (Fig. 3k). The *Nrxn3*_*Nlgn3* is well known for regulating astrocyte maturation as well as synapse development and function^41^. Thus, these findings strongly suggested a functional relay of the original astrocytes-GM to GM-relocated astrocytes-WM at the end of the acute phase of SCI.

To further confirm the injury-induced relocation of astrocytes-WM to the GM at 72 hpi, we performed immunostaining of the spinal astrocytes with anti-GFAP. The results not only verified the predominant localization and fibrous appearance of GFAP^+^-astrocytes in the WM of the intact and injured spinal cord but also showed a remarkable emergence of GFAP^+^-astrocytes in the GM after injury, and these GM-relocated GFAP^+^-astrocytes displayed a hypertrophic morphology (Fig. 3l). The loss of astrocytes-GM at 24 and 72 hpi and the striking occupation of massive GFAP^+^-astrocytes in the GM prompted us to further investigate the molecular origin and the potential functions of these GM-relocated astrocytes. To avoid confusion due to the discrepancy in the anatomic localization and the transcriptional features, we designated the injury-induced, GM-relocated astrocytes as “astrocytes-GMii” in the subsequent study.

### The astrocytes-GMii are molecularly distinct from the original astrocytes-WM/-GM and may contribute to spinal cord rescue/repair

We were keen to know where these astrocytes-GMii came from and what function(s) they played in the GM. Toward this end, we performed a series of exploratory data analyses and also cross-tested with the published scRNA-seq datasets of mouse SCI (Fig. 4a). As an attempt to address the question whether astrocytes-GMii were derived from local cell division or remote cell migration, we calculated and compared the proliferation score and the migration score (see Methods) at the spatial and cell type levels (Fig. 4b-e and Extended Data Fig. 12a, b). The results showed that significant cell proliferation was detected in the spots enriched by astrocytes-WM, reactive microglia and other cell types only at 72 hpi, much later than the time when a large number of astrocytes-GMii emerged. In contrast, the migration score started to increase in astrocytes-WM as early as 3 hpi, which kept rising at 24 hpi and expanded into the GM at 72 hpi. Notably, astrocytes-GM persistently showed the low proliferation and migration scores with a minimal increase of migration at 3 hpi. In addition, our analysis of the SCI scRNA-seq dataset^9^ also indicated a markedly higher migration ability associated with astrocytes-WM than astrocytes-GM (Extended Data Fig. 12b). Together, astrocytes-GMii likely originated from astrocytes-WM, which migrated into the GM and proliferated to compensate for the loss of original astrocytes-GM in the acute phase of SCI.

To reveal what unique molecular features astrocytes-GMii possess that made these cells distinct from the other astrocytes-WM and drove them into the GM upon injury, we assessed all the spots containing astrocytes-WM (astrocyte-WM spots) in our spatial transcriptomic dataset (Fig. 4f) using the anatomical definition by aforementioned UMAP clustering (Fig. 1g, k). In addition, to exclude potential confounding influences from other cell types in the same spot, only the 317 astrocyte marker genes used for cell type deconvolution were included. Despite considerable overlapping in their expression features that contribute to the general identity of astrocytes, these 317 marker genes could be clustered into four spatiotemporally-distinct astrocyte (SA) groups (Fig. 4g and Supplemental Table 5). Among them, the SA-1 group showed specifically high expression in the GM region at 24 and 72 hpi, while the SA-4 group also showed increased expression in the GM after injury, albeit to a lesser extent. We speculated that a SA score computed by these two SA gene sets would serve to spatially define the molecular features of astrocyte-GMii. Interestingly, the SA score was positively correlated with the proportion of remaining neurons in the spatial spots at 72 hpi (Fig. 4h, i). Together with the earlier SA-CCC analysis indicating a functional relay of astrocytes-GM to astrocytes-GMii at the end of the acute phase of SCI (Fig. 3j, k), these data suggested that the emergence of astrocytes-GMii and their interaction with neurons likely compensated for the loss of original astrocytes-GM and provided structural and nutritional support to the neurons spared from the primary injury.

To test the utility of the SA score for distinguishing astrocytes-GMii from the general population of astrocytes-WM in other spinal cord studies, we re-clustered the astrocytes-*Gfap* in the SCI scRNA-seq dataset^9^. Indeed, these cells were gated into two distinct populations based on the SA scores: astrocytes-GMii and the rest astrocytes-WM (-WMrs) (Extended Data Fig. 12c), which were also separated from each other in the UMAP plot (Fig. 4j). Next, to expand the spatially-defined molecular features of astrocytes-GMii, we identified the signature genes of astrocytes-GMii and -WMrs in the whole injured single-cell dataset (Extended Data Fig. 12d). Also, to facilitate precise sorting and genetic manipulation of astrocytes-GMii in the future, we further examined the potential surface markers and the upstream TFs of astrocytes-WMrs, -GMii and -GM (Extended Data Fig. 12e, f). It was worth noting that astrocytes-GMii were strongly associated with the increased activity of the Irf and Stat families of TFs, pointing to an immune-triggered transition of astrocytes-WM to astrocytes-GMii. Thus, the findings based on the spatially defined molecular features together the expanded signature genes of astrocytes-GMii demonstrated that our spatial transcriptomic analysis could provide valuable metrics to annotate independent cell states that are often hidden in common single-cell approaches.

To elucidate the molecular programming driving the transition of the original astrocytes-WM to the SCI-induced astrocytes-GMii, we performed pseudo-trajectory analysis of the SCI single-cell dataset by slingshot^42^. We identified continuous transitive structures between associated clusters in the low dimensional data (Fig. 4j, k) and highlighted the most dramatically altered genes along this transition (Fig. 4l). One of such genes was *Igfbp2* (*insulin-like growth factor binding protein 2*), and fluorescence ISH (FISH) of *Igfbp2* together with immunostaining of GFAP confirmed that astrocytes-GMii, represented by the *Igfbp2*^+^-GFAP^+^ double-positive cells, were specifically induced in the GM of injured spinal cord especially at 72 hpi (Fig. 4m). *Igfbp2* is a developmentally regulated gene that is highly expressed in the developing brain and markedly decreases after birth^43,44^, while its aberrant expression in cancer has been proposed to act as a hub of the oncogenic network that promotes tumor growth and metastasis^44,45^. Here, we showed that overexpression (OE) of *Igfbp2* in primary cultures of mouse astrocytes by lentiviral infection significantly promoted astrocyte migration in the *in vitro* scratch assay (Fig. 4n). Thus, the expression of *Igfbp2* not only can serve as a molecular marker for astrocytes-GMii but may also underlie the migration of astrocytes-WM to the GM in response to SCI.

Finally, to explicate the biological significance of astrocytes-GMii, we compared the DEGs and their enriched GO terms of astrocytes-WMrs, -GMii and -GM in the SCI single-cell dataset (Fig. 4o and Supplemental Table 6). The results indicated that astrocytes-GMii shared several neurotrophic functions in common with astrocytes-GM, which once again suggested a functional replacement of astrocytes-GM by astrocytes-GMii. In addition, the astrocyte-GMii signature genes such as *Vegfa, Gpr37l1, Fgfr3* and *Fgfr1* were associated with a protective role in neurological disorders^46^, which was consistent with the outcome of better neuronal survival observed in the higher SA scores of astrocytes-GMii (Fig. 4h, i). Together, as both the previous^47^ and this study showed that migrating astrocytes together with other cells in the SCI lesions expressed multiple axon/synapse-growth-supporting molecules, we anticipate that the surge of *Igfbp2*^+^-astrocytes to the GM of the injured spinal cord may provide neurotrophic support for spare neuron survival and spinal cord rescue in the acute phase of SCI (Fig. 4p). Future investigation is warranted for the finer classification of astrocytes-GMii and understanding of their spatial features and functional implications.

## Discussion

The spinal cord is a highly structured organ composed of various cell types that are well organized to perform complicated yet interdependent physiological functions. Two earlier spatial transcriptomic studies of the mammalian spinal cord identified pathway dynamics during the disease progression of ALS^13^ and described various cell transcriptional states over the course of injury-induced scar formation^48^. Meanwhile, several single-cell studies have demonstrated complex cellular heterogeneity and interactions in the mouse spinal cord^9,35,49-52^. How the diverse cell types respond and crosstalk to one another in a spatial- and context-dependent manner, especially in the early stage of SCI, drives the subsequent pathological process and is central to the pathophysiology of SCI. Here, we implemented the spatial transcriptomics to characterize the spatiotemporal programming of gene expression and cell organization in the immediate and the acute phase of SCI. We characterized and analyzed the spatiotemporal changes in gene expression, TF activities, regulon networks, cell compositions as well as cell-cell communications in SCI. To demonstrate the utility of the rich information generated in this study, we have set up an interactive web-portal for other researchers to explore *in situ* gene expression dynamics in SCI *(will release to the public upon acceptance of the paper)*.

Glial cells in the WM are generally considered to provide support and maintain extracellular homeostasis to neurons, forming myelin and clearing debris. Interestingly, our spatiotemporal transcriptomic data indicate that genes and cells in the WM respond to SCI more rapidly, robustly and sophisticatedly than those in the GM. The main molecular features of the WM shifted twice from Cluster 1 to 4 and then to 5, whereas the GM changed only once from Cluster 2 to 3 (Fig. 1). Moreover, the early response gene modules (e.g., CGM13 and CGM22), involving the MAPK/ERK signaling cascade, cytokine production and other immune and inflammatory responses, were activated in the immediate phase at 3 hpi in the WM but not the GM (Fig. 2). Thus, the glial cells in the WM do not merely insulate axons or respond subordinately to neuronal damage. Rather, they behave as the “first responders” in SCI and play a more active, leading and probably decisive role than previously thought.

SCI triggers multicellular gene expression changes in a temporally and spatially coordinated manner. When characterizing cell type-specific responses and cell-cell communications, single cell-based analyses are limited by the sampling bias in the tissue dissociation procedure and low-coverage sequencing^53^. To amend this, we took the advantage of spatial transcriptomics and developed a new computation strategy (SA-CCC) to unfold the cell communication by simultaneous evaluation of the physical distance between any cell-type pair and the locally aggregated expression level of a L_R pair that is supposed to express in this cell-type pair (Fig. 3). Using this approach, we not only systematically illustrate an overview of *in situ* cell-cell interactions of SCI, but also provide a useful toolset for evaluation of intercellular communications in a comprehensive multicellular context in biological processes and diseases.

Astrocytes are dynamic, diverse cells whose molecular and functional heterogeneity is closely related with the local environment^54,55^. Despite their being a barrier to axonal regeneration in the later phases, migrating reactive astrocytes that form the glial scar play a beneficial role by the closure of the wound and segregation of immune/inflammatory cells, which contribute to the spontaneous, partial functional recovery in the subacute phase^56,57^. And, cell-based treatments including the induction of differentiation and migration of reactive astrocytes have emerged as a potential therapeutic strategy to improve the functional recovery in patients^58-61^. However, the specific dynamics and the regulatory programs of astrocyte migration in SCI are still poorly understood, largely due to the difficulty in exploring these mechanisms *in vivo*. In this study, by leveraging the spatiotemporal transcriptomic allocation and profiling different cell types and subtypes, we uncover a distinct population of injury-induced reactive astrocytes (astrocytes-GMii) and explicitly show that these astrocytes originate from the WM and migrate to the GM, in which they interact with spare neurons and other cell types at 24 and 72 hpi. Thus, unlike the double-edged scar-forming astrocytes and those promote detrimental secondary inflammation and immune responses in the subacute and the intermediate phases, this population of astrocytes-GMii respond much rapidly in SCI and function as a hub of the multicellular communication and re-organization in the acute phase, which may support neuronal survival and spinal cord rescue after the primary injury. In addition, the pseudotime trajectory and the DEG analysis strongly suggested that the local environment of the GM triggered the transition of the relocated astrocytes-WM to astrocytes-GMii, and the latter shared certain gene expression similarity with the original astrocytes-GM, especially regarding the neurotrophic function in maintaining neuronal synapses. Nevertheless, these astrocytes-GMii do not necessarily migrate from the same segmental level of the injured spinal cord^62^ and are unlikely through tangential astrocyte migration that is absent in adult spinal cord even after injury^63^.

Astrocytic scars have been studied for several decades; however, the exact molecular mechanisms regulating their formation are not fully understood. This is in part due to the lack of individual markers for the diverse range of subtype- and context-specific astrocytes. For example, in this study, GFAP is strongly expressed in both the original astrocytes-WM and the SCI-induced astrocytes-GMii, despite that their spatial distributions, morphologies, molecular features and cellular functions are quite different. By a series of integrated spatial and single-cell transcriptomic analyses, we not only demonstrate that astrocytes-GMii are molecularly distinct from the original astrocytes-WM and -GM but also identify that these astrocytes-GMii express both the reactive astrocyte marker GFAP and the IGF signaling factor IGFBP2, and the latter may underlie the SCI-induced migration of astrocytes-WM to the GM. Hence, our work exemplifies how the spatiotemporal transcriptomic signatures combined with single-cell resolution may facilitate subtype annotation and functional interrogation in SCI as well as other physiological and pathological conditions. In addition, although the secreted protein Igfbp2 was believed to play an essential role in promoting neurite outgrowth in developing and injured brains^64,65^, a recent study showed that blocking Igfbp2 in the astrocyte-conditioned medium derived from a Rett syndrome model induced a significant increase in neurite outgrowth^66^. Thus, future efforts are warranted to understand the specific role of the increased expression *Igfbp2* in these astrocytes-GMii and to induce the astrocytes-GMii to adopt a neuroprotective and pro-regenerative state that may aid spinal cord repair.

Taken together, our work here provides the most comprehensive gene expression profiling of SCI (to the best of our knowledge), involving injury time, distance, directionality, anatomic domain, cell type and other dimensions. As summarized in Extended Data Fig. 13, this study disentangles the SCI-induced complex transcriptional programs and multicellular responses in the tissue context and establishes a spatiotemporal transcriptomic landscape for in-depth research in the future, providing an open data resource and a platform for identifying potential targets for therapeutic intervention of SCI.

## METHODS

### Animal Care

All practices on mice in this study were performed in compliance with the institutional guidelines on the scientific use of living animals at the Shanghai Interdisciplinary Research Center on Biology and Chemistry, the Chinese Academy of Sciences (CAS). Animal distress and conditions requiring euthanasia were addressed and the number of animals used was minimized. All mice were housed in a pathogen-free barrier facility with free access to food and water.

### Mouse spinal cord injury (SCI)

The C57BL/6J mice (Jackson Laboratory, #000664) used in this study were purchased from the Shanghai Laboratory Animal Center (SLAC). Adult female mice of 8-9 weeks old were used in the SCI. The spinal cord transection was performed as previously described^67^. Briefly, after exposure and removal of the lamina of the tenth thoracic vertebra (T10) spinal segment, the spinal cord was transected with Micro Scissors, and the wound was then cleaned and sutured. For sham controls, only laminectomy at T10 was performed. For post-operative care, the mice received analgesics at the wound every 12 h and were subjected to manual bladder-emptying at least twice daily.

### Behavioral evaluation of SCI

All animals were accommodated to the testing room or apparatus for at least 1 h before behavioral assessments. Blind scoring was performed to ensure that observers were unaware of treatments. The animals were evaluated for the hindlimb locomotion performance at 0 (uninjured), 3, 24 and 72 hours post injury (hpi) by the Basso mouse scale (BMS)^68^. Briefly, mice were placed in an open field for 4 min. Two observers assessed the performance independently with the reference to the motion attributes of the hindlimbs, including joint movement, weight support, plantar stepping, coordination, paw position, and trunk and tail control. The BMS scores range from 0 to 9 points (0 means complete paralysis and 9 means normal mobility). For the mechanical sensitivity (von Frey) test^69^, the mice were placed on the bottom of a metal grid and the mid-plantar surface of the hindpaw of the mice was stimulated with 2 g von Frey filaments vertically. The filaments were applied to the hindpaw surface until the filament bent, and paw withdrawal or flinching was considered as a positive response. The test of tactile withdrawal threshold was repeated 10 times in each mouse. The mean value of the performance of the left and right hindlimbs was calculated and used in the BMS and von Frey tests.

### Cryosection and immunohistochemistry

Anesthetized mice were fixed via transcardial perfusion with diethylpyrocarbonate-treated phosphate-buffered saline (PBS) followed by 4% (w/v) paraformaldehyde (PFA, RNase-free; Bioss, C2055). Spinal cord tissues were then dissected and post-fixed in 4% PFA at 4°C overnight. The fixed spinal cord samples were cryoprotected in 30% (w/v) sucrose (Sangon, A0498) in PBS, embedded in Tissue-Tek O.C.T. (Sakura, 4583), snap-frozen in liquid nitrogen, and stored at -80°C until cryosection. Before cryosection, samples were equilibrated to -20°C in the cryostat for at least 30 min. Cross sections (20 μm) of the spinal cord were collected on the slides and could be stored at -80°C up to 2 weeks.

For hematoxylin and eosin (H&E) staining, the spinal cord sections on the slides were incubated at 37 °C for 1 min,and then completely immersed in the prechilled methanol at -20 °C for 30 min. After three washes with deionized water (diH_2_O), isopropanol was added to fix the tissue slides at room temperature (RT) for 1 min. Thereafter, the slides were air-dried and hematoxylin (Agilent, S330930-2) was added to the slides at RT for 7 min. After three washes with diH_2_O, slides were treated by Bluing Buffer (Agilent, CS70230-2) at RT for 2 min. Finally, eosin (Sigma, HT110216-500ML) was added at RT for 1 min and the slides were washed in diH_2_O for three times, followed by imaging with Aperio LV1 slide scanner (Leica).

For immunostaining, the spinal cord sections on the slides were incubated at 37 °C for 1 min, post-fixed once more in 4% PFA at RT for 15 min, and then permeabilized and blocked with the blocking buffer (10% normal goat serum and 0.5% Triton X-100 in PBS) at RT for 1 h. The primary rat anti-GFAP antibody (Invitrogen, 13-0300) was subsequently included in the blocking buffer at 4 °C overnight. The sections were washed with 0.3% Triton X-100 in PBS for three times and the secondary goat anti-rat-Alexa Fluor 568 antibody (Life Technologies, A11077) was added to the slide for 1 h at RT. After washed by 0.3% Triton X-100 in PBS for three times, the spinal cord sections on the slides were mounted with coverslips using the VECTASHIELD Antifade Mounting Medium with DAPI (Vector Laboratories, H1200) and imaged by Dragonfly Spinning Disk Confocal Microscope (Andor) with a 20x/0.75 NA objective.

### *In situ* hybridization (ISH) and fluorescence *in situ* hybridization (FISH)

ISH was performed as previously described^70^. Briefly, the mouse spinal cord sections were treated with 10 μg/mL proteinase K (QIAGEN, 19131) at 37°C for 5 min and then post-fixed in 4% PFA at RT for 10 min. After washing and dehydration, DIG-labeled RNA probes at approximately 1 μg/mL in hybridization buffer were incubated with the sections on the slides at 65°C overnight. Afterwards, the slides were washed with 50% formamide (Sangon, A600212) in 2x saline sodium citrate (SSC) (Invitrogen, AM9770) at 65°C for 30 min, followed by incubation with 1 mg/mL Rnase A (Roche, 10109142001) at 37°C for 30 min. The slides were then washed sequentially in 2x SSC at 60°C for 20 min, 0.2x SSC at 60°C for 20 min, and 0.1x SSC at RT for 20 min. For ISH, the slides were incubated with anti-DIG-AP (Roche, 11093274910) at 4°C overnight, followed by washing and incubation with BM-purple (Roche, 11442074001) until dark purple color became visible.

The composition of the hybridization buffer: 50% (v/v) formamide; 10 mM Tris-HCl solution, pH 8.0 (Sangon, B548127); 200 μg/mL yeast tRNA (ThermoFisher, AM7119); 10% (v/v) dextran sulfate (Millipore, S4030); 1x Denhardt’s solution (Sigma, D9905); 600 mM NaCl, RNase-free (ThermoFisher, AM9760G); 0.25% SDS (ThermoFisher, AM9820); 1 mM EDTA (0.5 M), pH 8.0, RNase-free (ThermoFisher, AM9261); Nuclease-free water (ThermoFisher, AM9937).

For FISH, the procedure was similar to the ISH but the anti-DIG-POD antibody (Roche, 11207733910) was used for signal development. The slides were finally incubated with TSA_Plus_Fluorescence Kits (Akoya, NEL745001KT) according to the manufacturer’s instruction and imaged.

For simultaneous detection of the immunofluorescence signals of GFAP with FISH, the immunostaining was conducted after FISH images had been obtained. Antigen retrieval was performed with Antigen Retrieval Solution (Solarbio, C1035) before the conventional immunostaining procedure.

The RNA probes used in this study were adapted from the Allen Spinal Cord Atlas (https://mousespinal.brain-map.org/), and synthesized with the DIG RNA labeling kit (Roche, 11277073910) following the procedure described in (Peng et al., 2016)^70^. The following primers were used:

*Gfap* forward: 5’---GTGGATTTGGAGAGAAAGGTTG---3’

*Gfap* reverse: 5’---CTGGAGGTTGGAGAAAGTCTGT---3’

*Pla2g7* forward: 5’---AAGGTCGCCTCGACACTG---3’

*Pla2g7* reverse: 5’---TCAAAGGGTGACCCAGGA---3’

*Igfbp2* forward: 5’---ACAGTGATGACGACCACTCTGA---3’

*Igfbp2* reverse: 5’---CCTCTCTAACAGAAGCAAGGGA---3’

### Culture primary astrocytes from mouse pup brains

Primary astrocytes were isolated from C57BL/6J newborn mice at postnatal day 2 (P2) as previously described^71^. In brief, after removal of the meninges, mouse cortical tissues were minced and digested sequentially with 0.05% DNase I (Sigma, DN-25) for 5 min and 0.05% trypsin (Gibco, 15090-046) for 20 min at RT, followed by filtration through a 70-μm cell strainer (FALCON, 352350) to remove undissociated tissues and debris. The filtrate was centrifuged at 150 *g* for 10 min, and the cell pellet was resuspended after decanting the supernatant. The harvested cells were plated into T-75 flasks coated with 50 μg/mL poly-L-Lysine, and incubated at 37 °C with 5% CO_2_ in an incubator. The growth medium (10% fetal bovine serum and 2% penicillin-streptomycin in DMEM) was changed every other day. Upon the confluence of the cells, oligodendrocyte progenitor cells (OPCs) could be detached by tapping the flask and DNase I digestion (37 °C for 5 min). After completely removing the supernatant, the remaining adherent cells were subsequently digested with 0.25% trypsin for 5 min. The cell suspension was mixed with an equal volume of the growth medium to deactivate trypsin, and centrifuged for collecting the pellet. Thereafter, the resuspended cells were subjected to two rounds of incubation in bacterial grade plates for 20 and 90 min, respectively. This helped to effectively remove the microglia as they are more prone to attach to the plate surface compared with astrocytes. The supernatant was then transferred to a new T-75 flask, and primary astrocytes were allowed to grow to confluence before use.

### Lentivirus production

To generate lentivirus for infecting primary astrocytes, 293T cells were co-transfected with target plasmid (pCDH-vector-flag or pCDH-*Igfbp2*-flag), psPAX2 and pMD2.G with a ratio of 4:3:1 in Opti-MEM (Gibco, 31985070) using Lipofectamine 2000 (Invitrogen, 11668). The culture medium was collected at 48 h after transfection and passed through a 0.45-μm filter. Viral particles were concentrated using the Lenti-X Concentrator (Clontech, 631232) and the viral pellets were resuspended in PBS for astrocyte infection.

### The *in vitro* cell migration assay

Primary cultures of mouse astrocytes were grown *in vitro* in the plates coated with 5 μg/mL fibronectin (Sigma, F2006) for 4-5 days before they were infected with lentivirus. Three days after infection, the astrocytes were treated with 10 μg/mL mitomycin C (Sigma, M5353) for 2 h to suppress cell proliferation and then the middle of the monolayered astrocytes was scratched with a pipette tip to create a gap (cell-free area), which was traced and imaged at 0, 24 and 48 hours post scratch (hps) using phase contrast microscopy.

### Image acquisition, processing and quantification of light microscopy

Brightfield images of H&E staining and ISH were acquired using the Aperio LV1 slide scanner (Leica) and processed in ImageScope. FISH and immunofluorescence images were acquired using the Dragonfly Spinning Disk Confocal Microscope (Andor) with a 20x/0.75 NA objective. The 3D rendering of the GFAP^+^-*Igfbp2*^+^ astrocytes was performed in Imaris using the Surface function. The images were processed and assembled into figures using Adobe Photoshop 2022 and Adobe Illustrator 2022.

For quantification of astrocytes, two regions of interest (ROIs) were drawn to outline the left and right grey matter (GM) of each spinal cord section from three different animals (n = 6 ROIs for each time point). The number of DAPI, GFAP^+^ or *Igfbp2*^+^ cells within each ROI was counted by ImageJ using the Analyze Particles function. And the GFAP^+^/*Igfbp2*^+^ double-positive cells over DAPI were visually examined and manually counted. Statistical analysis was determined by one-way analysis of variance (ANOVA) with Tukey’s HSD post-hoc test at \**p* < 0.05, \*\**p* < 0.01, and \*\*\**p* < 0.001. Error bars represent the standard error of the mean (SEM).

For quantification of the gap size in the cell migration assay, the cell-free area was delineated, with its size measured by ImageJ at 0, 24 and 48 hps. Six randomly-selected, non-overlapping regions of the samples prepared independently in each group were included at each time point for calculating the percentage of the gap area, using the following equation: Relative gap size (%) = S/S_0hps_ x 100%, where S stands for the number of the pixels of the cell-free area of a sample, and S_0hps_ is that of the same sample at 0 hps.

### Acquisition of mouse spinal cord sections for 10x Genomics Visium spatial RNA-seq

Dissected spinal cords were washed with 1x PBS and subsequently embedded in Tissue-Tek O.C.T. through a bath of dry ice and prechilled ethanol. Before sectioning, Visium slides and samples were equilibrated to -20°C in the cryostat for at least 30 min. Four consecutive spinal cord cryosections were cut at 20-μm thickness each and attached to one of the four capture areas of the Visium Spatial Gene Expression slides (10x Genomics, 1000184). The slides were stored individually in sealed containers at -80°C before library construction.

### Spatial transcriptomics library construction and sequencing

The spatial transcriptomics library was constructed according to the Visium Spatial Gene Expression Reagent Kits User Guide (10x Genomics, CG000239 Rev D). In brief, after H&E staining and imaging, the spinal cord samples on the slides were permeabilized for 18 min, which was determined based on the pre-tests using the tissue optimization kit (10x Genomics, 1000193). Thereafter, the tissue slides were reverse transcribed and amplified, producing cDNAs containing the spatial barcodes. The amplified cDNAs were subjected to library construction and sequenced with the Illumina NovaSeq 6000 system using a 150-bp paired-end setting at a depth of 100k reads per spot.

### Data analyses of spatial transcriptomics

#### Pre-processing and quality control

The raw Visium spatial RNA-seq data and histological H&E images were processed using the Space Ranger pipeline (v.1.0.0, 10x Genomics) to align and summarize unique molecular identifier (UMI) counts according to the reference genome “Mouse Genome mm10”.

To assess the quality of the spatial RNA-seq data, we grouped the SCI samples based on the replicate batch and estimate the sample correlation coefficients. In brief, average expression of each spinal cord section replicate was calculated and the Pearson correlation coefficient (PCC) analysis was performed across all samples. The expressed genes, UMI counts and mitochondrial levels were computed by Seurat (version 4.1.1)^72^. Gene expression levels of all spatial spots were normalized and scaled using the ‘SCTransform’ function^73^ and merged by regressing out ‘section’ factors to create a single Seurat object for further analyses. All the hemoglobin-related genes (due to infiltration of red blood cells from the damaged blood-spinal cord barrier) were removed from the matrix before merging. Other potential confounding effects including cell cycle, sequencing library size and percentage of mitochondria were regressed out as well.

#### Spatial cluster analysis

The top 3,000 highly variable genes during SCI were identified and used for the subsequent analysis. We computed 20 principal components (PCs) with ‘runPCA’ and constructed the shared nearest neighborhood graph with 50 local neighbors of each pixel using the ‘FindNeighbors’ function. We used the Louvain clustering algorithm to optimize the modularity of neighbors and the UMAP (Uniform Manifold Approximation and Projection) visualization at the resolution of 0.25. Seven spatiotemporal clusters were recognized and classified into four anatomic domains according to their spatial distributions and the identified signature genes: white matter (WM), middle grey (MG), dorsal horn (DH) and ventral horn (VH).

#### Evaluation of activity scores of gene sets

Injury, proliferation and migration scores of the indicated gene sets were computed using the function of ‘AddModuleScore’ in Seurat with gene sets extracted from published literatures or defined by the enrichment analysis. Specifically, the injury score was calculated based on the expression of the following 22 genes: *Adamts1, Atf3, Ccl2, Ccnd1, Cd68, Cebpd, Cyba, Fn1, Gal, Gap43, Hmox1, Hspb1, Igfbp2, Jun, Junb, Fos, Lgals1, Neat1, Socs3, Tnc, S100a10*, and *Timp1*; the proliferation score was calculated based on the expression of *Mki67*; and the migration score was calculated based on the expression of the genes in the “positive regulation of cell migration” gene ontology (GO) term.

#### Differential expression gene (DEG) and GO term analyses

For DEG analysis, ‘FindAllMarkers’ and ‘FindMarkers’ of Seurat were run with the following parameters: logfc.threshold = 0.5, min.pct = 0.25 and method = “wilcoxon”. GO term analysis was done using the “clusterProfiler” R package^74^. The ‘compareCluster’ function was used for gene list enrichment against all expressed genes by setting *p*. adjust < 0.01 or 0.05. The top 5 or 10 (*p*. adjust) terms were displayed by DotPlot.

#### Gene co-expression analysis

The WGCNA (weighted gene co-expression network analysis) package in R^75^ was used to measure co-expression networks across all the spatial spots. Genes expressed in over 100 spatial spots were selected and their average expression levels in each group of injury time and anatomic domains were calculated. To calculate the adjacent matrix, the soft power of 18 was chosen with the WGCNA function ‘pickSoftThreshold’. The module identification and topological connectivity network were computed using the ‘DynamicTreeCut’ function with min.module.size = 30, deep = 2 and mergeHeight = 0.15. The most connected genes in the topological network of each module were defined as the hub genes and were visualized with Cytoscape^76^. Ribosome-related genes were filtered manually to avoid overdominance in co-expression analysis when displaying connectivity network of Co-expressed Gene Module (CGM) 9. Enriched regulons of CGMs were determined using the hypergeometric test against previously identified regulons of transcription factors (TFs) and the cutoff *p*. adjust (which was calculated with the bonferroni multiple comparison correction of *p* value in hypergeometric test by the number of modules) was set to < 0.05. Indices of significance were calculated by (0.05 -*p*. adjust)/0.05. The top 3 (ranked by significance index) regulons of gene modules were displayed.

#### Analysis of transcriptional regulons and regulon modules (RMs) with SCENIC (single-cell regulatory network inference and clustering)

The Python package “pyscenic”^28^ was used to detect active transcription RMs. Briefly, the spatial gene expression matrix was first filtered to exclude genes expressed in fewer than 0.1% of all the spots. The gene-gene correlation matrix for module identification was computed with the Random Forest based on the ‘GRNBoost2’ algorithm. Each module was further pruned to only include genes that were present in the RcisTarget database with the ‘prune2df’ function. Therefore, regulons consisting of the TFs and their transcriptional targets in the co-expression module were detected finally. Overall, 515 regulons were found in this study and the regulon scores were calculated by the ‘AUCell’ function. To evaluate the co-expression relationship of the regulons, the top 20 regulons (based on the regulon specificity score, the RSS^77^ for each combination of anatomic domain and injury time was picked (147 regulons in total; less than predicted due to redundant regulons), which were then used to compute their connectivity specificity index (CSI) for each pair of regulons following the instruction in (Bass et al., 2013)^78^. Based on the CSI, 7 RMs were identified via hierarchical clustering. Activity score of each CSI module was calculated by averaging the regulon scores within each module. Regulon modules with PCC > 0.65 were visualized with Cytoscape^76^.

For regulon analysis of the single-cell data of astrocytes, the same procedure was followed and 433 regulons were identified. The scaled average activity was computed across three spatial associated states and the top 10 regulons of each state (based on RSS) were shown.

#### Cell type deconvolution and identification of cell type-enriched DEGs

To estimate the proportions of the cell types in each spatial spot, all the spots were deconvoluted with the “Spatial eXpression R (spacexr)” package (version 2.0, formerly RCTD^34^). For a better inclusiveness of diverse cell types, we imported and combined two mouse spinal cord single-cell datasets^9,35^ in the RCTD analysis. We further defined astrocytes in the SCI single-cell data^9^ into three subtypes (astrocyte-*Gfap*, -*Slc7a10* and -*Svep1*) with the annotation by Rosenberg et al. (2018)^35^ as the reference by label transfer^79^. The cell types in the sham and injured spinal cords (up to 72 hpi) identified by Milich et al. (2021)^9^ were used to construct the spacexr reference. Finally, the percentage of different cell types for each spot was calculated using ‘run.RCTD’ with “multi” mode (CELL_MIN_INSTANCE = 10, UMI_min = (min(puck@UMI)-1), UMI_max = (max(puck@UMI)+1), fc_cutoff_reg = 1). For robust cell type decomposition, confident sub_weight was used to evaluate the final proportions of different cell types in all spots. Astrocytes-*Svep1* and dendritic cells were merged into the class of “other cells” due to their extremely low presence in the overall spatial transcriptomic data and were not considered in the subsequent analysis. In addition, excessive subtype classifications showing scattered spatial distributions were merged and analyzed at the cell type level.

The DEGs of the indicated cell type were calculated with the ‘C-SIDE’ function^80^. For analysis of astrocyte subtype-enriched DEGs, astrocytes-WM or -GM in the four replicates of the +1.0 mm sections at 0 hpi were passed to C-SIDE with the parameters of “cell_type_threshold = 1, weight_threshold = 0.9 and doublet_mode = F” and their spatial localization was used as a function of the covariates.

#### Spatial-aware cell-cell communication (SA-CCC) analysis

The ligand_receptor (L_R) pair interaction information was extracted from CellChatDB.mouse of CellChat^37^. To further pinpoint the L_R pair interactions between different cell-type pairs in the spatial context, we binarized the L_R pair consensus matrix of each spatial spot based on whether its value exceeded its assembly mean across all spots. Similarly, the cell-type pairs were also binarized depending on their deconvolution proportions. To minimize the interference from ubiquitous and/or ambiguous interactions from other cell types, the proportions of neurons and oligodendrocytes were filtered by their mean values when assessing the spatial interactions. Additionally, to exclude “false interaction” between spatially nonadjacent cells, the cell-type pairs were eliminated if the Spearman rank correlation coefficient of their deconvolution proportions was < -0.4. The spatial correlation between each L_R pair and cell-type pair was calculated using the one-side Fisher’s exact test. When the *p* value was < 0.001, the cell-type pair was considered to have a bona fide spatial cell-cell interaction. The expression level of a specific L_R pair for an associated cell-type pair was calculated by averaging the values of L_R pair geometric mean (GeoM) across their colocalized pixels. The aggregated types of cell-cell interactions, the average expression levels of all L_R pairs between indicated cell-type pairs and the -log10(GeoM of *p* values) at each time point of SCI were visualized by igraph^81^. Only interactions between the cell-type pairs that have over 10 shared positive spatial spots were shown. The SA-CCC results were further applied to assess the cell-cell interactions identified in the SCI single-cell study^9^ using the CellChat with the default parameters. The SA-CCC analysis is accessible and searchable in our interactive data exploration portal: *(will release to the public upon acceptance of the paper)*.

#### Re-analysis of the SCI single-cell datasets of the spatially-defined astrocytes

To re-cluster the astrocytes in the SCI single-cell dataset^9^, the single-cell matrix was normalized and scaled using the ‘SCTransform’ function (var.to.regress = c(“orig.ident”,”S.Score”,”G2M.Score”,”percent_mt”,”percent_rp”), do.scale = TRUE). Cells with low counts (< 2,500) were filtered. Ribosome- and mitochondria-related genes were regressed out. The top 16 PCs, 40 k.param and resolution = 0.2 were used in the UMAP analysis (Extended Data Fig. 9d). Two molecularly-defined subtypes of astrocytes-WM (*Gfap*^+^) and astrocytes-GM (*Slc7a10*^+^) were identified, and the subsets of the RNA matrix of the astrocytes were used in the subsequent analyses.

The spatially scaled data were used to extract the expression profiles of astrocytes-WM in all the spots that contained astrocytes-WM based on the deconvolution result. The 317 astrocyte marker genes used in the cell type deconvolution analysis were retained. Scaled expression was binarized into 1 and -1 based on the threshold = 0, which was used to estimate the expression of each marker gene at the bulk level in the WM and the GM at all time points of SCI. K-means clustering (k = 4) was employed to identify spatially-defined marker genes for the SA groups of astrocytes in Fig. 4g.

To evaluate whether the SA-1 and SA-4 gene groups were able to distinguish astrocytes-GMii from the rest astrocytes-WM, the astrocytes-*Gfap* (WM) defined in the SCI single-cell study^9^ were re-clustered using the maximum distance and the ‘cutree’ function based on the scores of the SA-1 and SA-4 gene sets. The DEGs of the original astrocytes-WM and -GM as well as the injury-induced astrocytes-GMii were identified and compared pairwise to reveal their overlapping relationships (logfc.threshold = 0.5, *p*. adjust < 0.01). Surface marker analysis was done by overlapping the surface genes downloaded from Cell Surface Protein Atlas with astrocyte-GMii signature genes.

#### Pseudotime analysis

Trajectory inference was performed in low dimension data of the astrocytes-*Gfap* (WM), including the astrocytes in the uninjured WM and the GM-relocated, WM-featured astrocytes after injury, in the SCI single-cell study^9^ using the slingshot package (v2.5.5)^42^.

#### Statistical information

The exact statistical information of the comprehensive spatial transcriptomic analyses in this study is specified in each related section of the Methods above or as indicated in the legends of the relevant figures.

## Data and Materials Availability

Raw data generated by SpaceRanger pipelines are available through the NODE project under the accession number OEP003508. Our SCI spatial transcriptomic resources can be explored at the SCI-ST web portal *(will release to the public upon acceptance of the paper)*. All other data are available in the main text and the supplementary materials. All unique and stable reagents generated in this study are available from the corresponding authors with a completed Material Transfer Agreement (MTA).

## Supplemental Information

Supplemental information includes Extended Data Fig. 1-13, Supplemental References 82-86, and Supplemental Tables 1-6.

## Acknowledgements

We thank Drs. J. Hu, Z. He, C. Zhang, and Misses. J Zhu and Z. Sun for assistance in mouse operation, Dr. Q. Wang for help with cryosection, Mr. G. Zhang for technical support with 10x Genomics Visium spatial RNA-seq, Dr. X. Sun for help with ISH, and Drs. W. Li, Y. Song, B. Zhou, P. Yu, C. Liu, Z. He, G. Bai, H. Liu and J. Yuan for comments and critical reading of the manuscript.

## Author Contributions

AL, GP and YF conceived the research; ZW, SS, and YF designed the experiments; ZW, SS and TL performed the experiments and analyzed the data; AL, WY and YL contributed important reagents; ZL, JX and GP performed the bioinformatic analyses; ZW, ZL, TL, GC, KZ, GP, AL and YF interpreted the results; ZW, ZL, TL, GP and YF prepared the figures; ZW, ZL, TL, AL, GP and YF wrote the paper. All authors read and approved the final manuscript.

