## Extended Data for "Spatiotemporal orchestration of multicellular transcriptional programs and communications in the early stage of spinal cord injury"

#### Supplementary Inventory

1. **Extended Data Fig. 1 to 13**

2. **Supplemental References 82-86**

3. **Supplemental Tables 1 to 6 (in separate spreadsheets)**

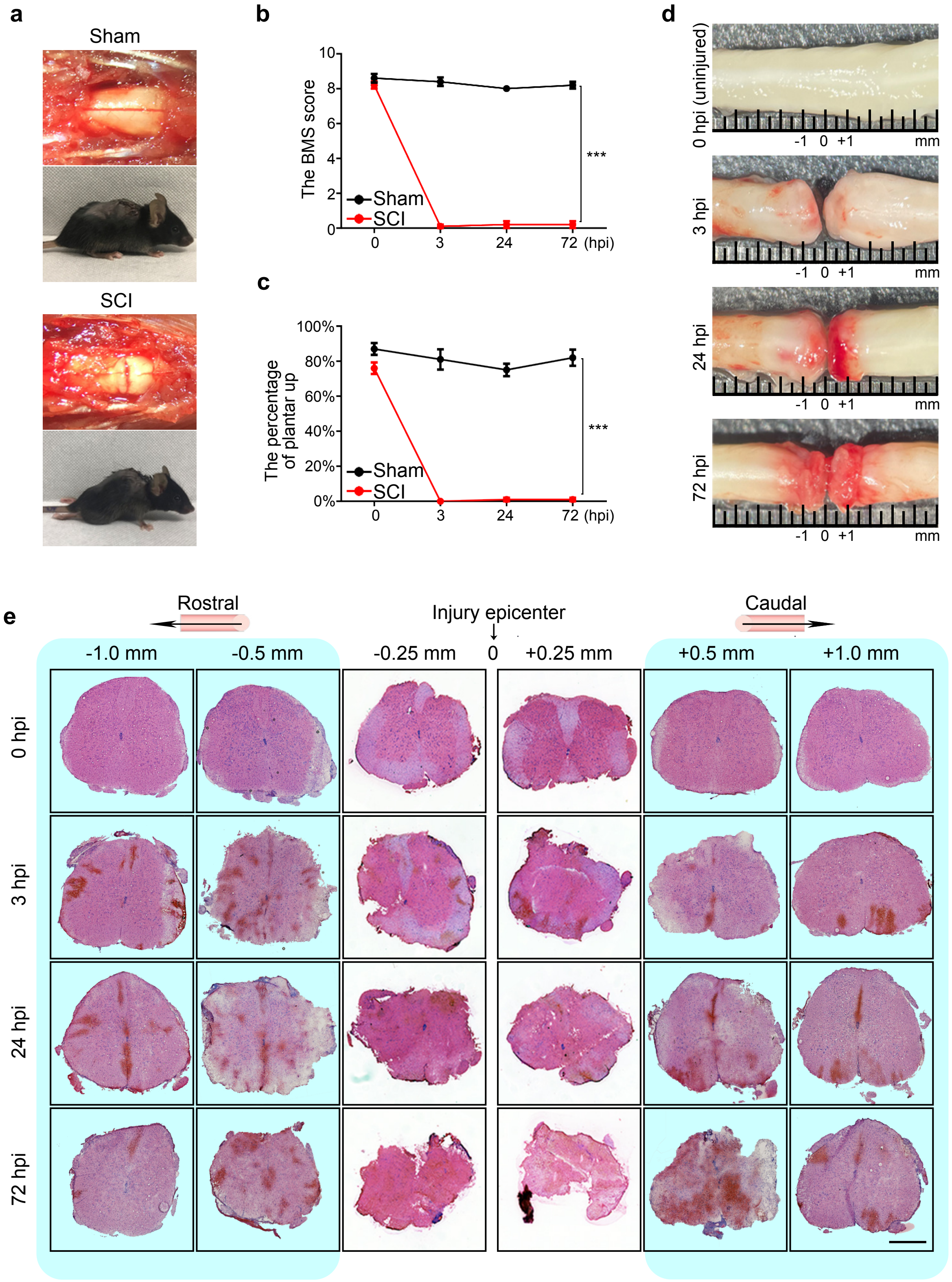
a
Sham
SCI
b
The BMS score
10
8
6
4
2
0
Sham
SCI
0
3
24
72 (hpi)
***
c
The percentage of plantar up
100%
80%
60%
40%
20%
0%
Sham
SCI
0
3
24
72 (hpi)
***
d
0 hpi (uninjured)
3 hpi
24 hpi
72 hpi
-1 0 +1
mm
e
Rostral
Injury epicenter
Caudal
-1.0 mm
-0.5 mm
-0.25 mm
0
+0.25 mm
+0.5 mm
+1.0 mm
0 hpi
3 hpi
24 hpi
72 hpi

**Extended Data Fig. 1. The setup and examination of the mouse SCI model**

(**a**) Representative photographs of the exposed mouse spinal cord at T10 and the mice showing the ability of the hindlimbs to support the body after sham (upper panels) or transection operation (lower panels).

(**b, c**) Behavioral evaluation of the hindlimb functions of the sham or SCI group of mice by the BMS score (**b**) or the von Frey test (**c**) at 0 (uninjured), 3, 24 and 72 hours post injury (hpi). Data are shown as mean ± SEM; n = 5; ****p* < 0.001; two-way ANOVA.

(**d**) Photographs showing the gross morphology of the mouse spinal cord at the indicated time points of SCI.

(**e**) The H&E staining of the spinal cord sections (20-μm thickness) at the indicated distances, directions and time points of SCI. Note a severe deformation at 0.25 mm from the injury site. As such, only the sections at the distances > 0.25 mm (highlighted in blue) are included in the spatial transcriptomic analyses in this study.

**Extended Data Fig. 2 - SCI ST**

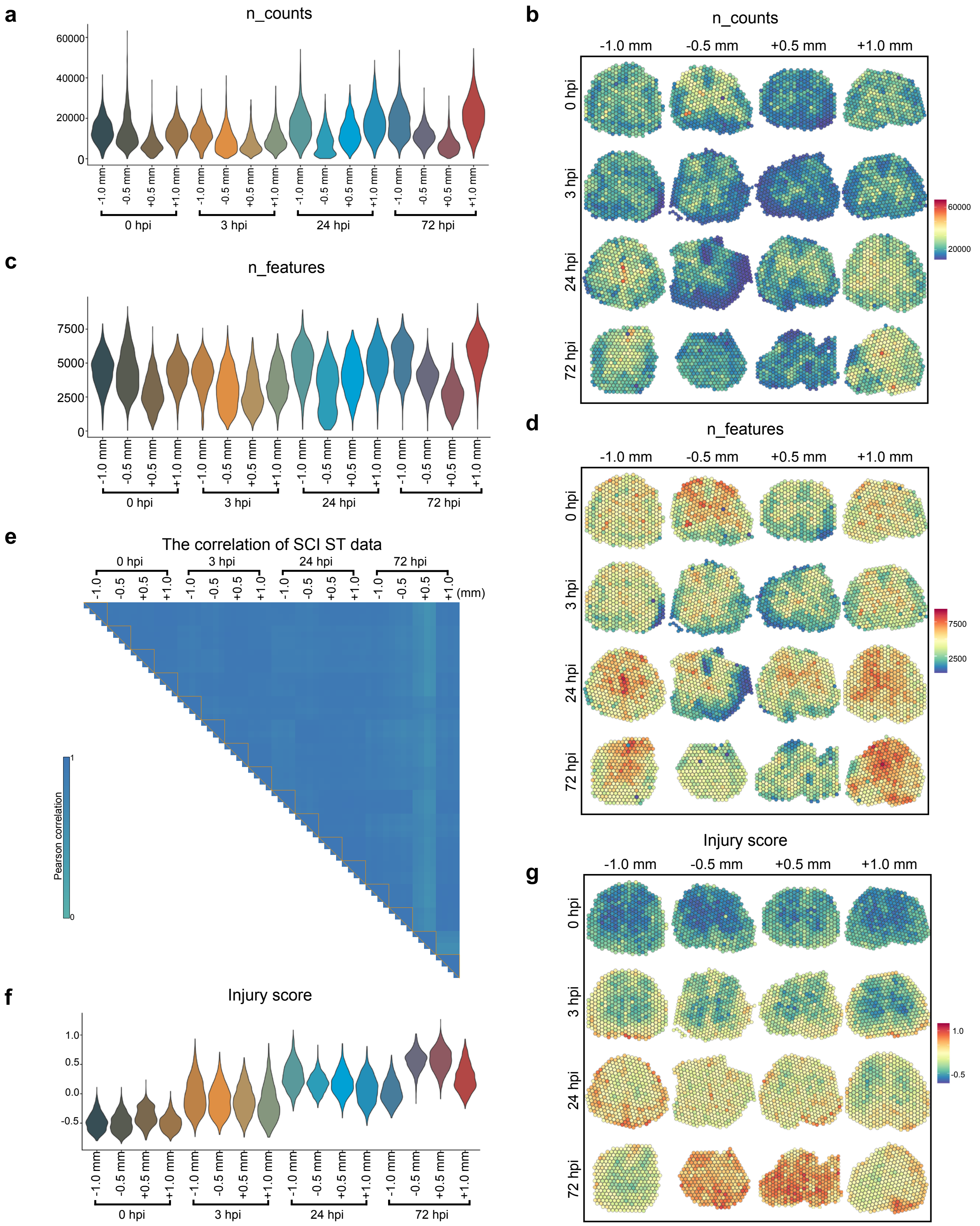

**Extended Data Fig. 2. Quality control of the spatial transcriptomic data**

(**a-d**) The total read counts (**a**, **b**) or the numbers of detected genes (**c**, **d**) of the 4 replicates of the indicated spinal cord sections at the specified time points of SCI in the spatial RNA-seq are shown in the violin (**a**, **c**) and spatial (**b**, **d)** plots.

(**e**) Pearson correlation coefficients of the bulk gene expression levels of all the 64 SCI samples in this spatial transcriptomic (ST) study.

(**f**) The scaled injury scores of the 4 replicates of the indicated spinal card sections at the specified time points of SCI are shown in violin plots.

(**g**) The spatial view of the injury scores of the indicated spinal cord sections at the specified time points of SCI are shown.

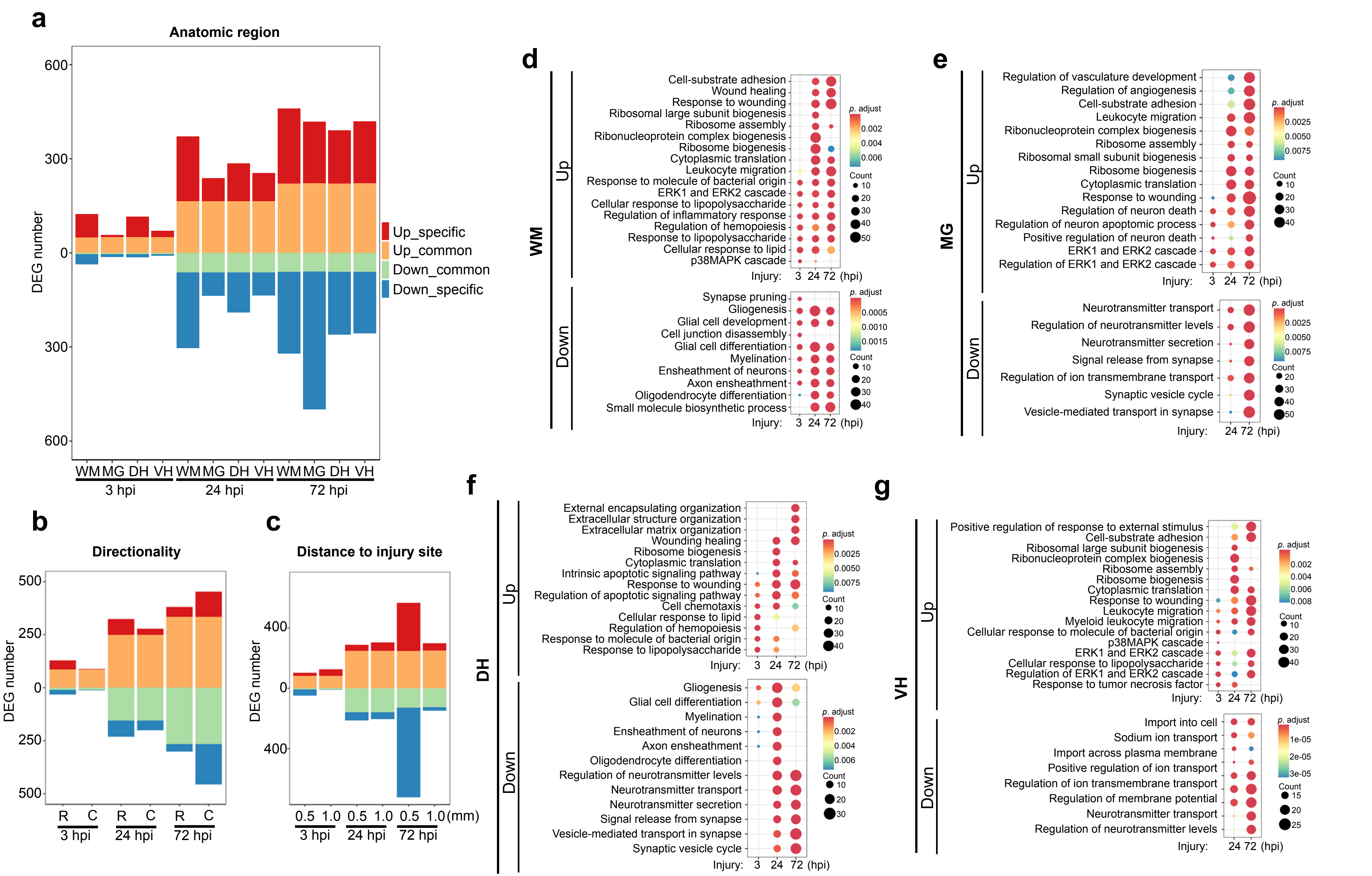
a
Anatomic region
DEG number
Up_specific
Up_common
Down_common
Down_specific
WM MG DH VH WM MG DH VH WM MG DH VH
3 hpi
24 hpi
72 hpi
b
Directionality
R C R C R C
c
Distance to injury site
0.5 1.0 0.5 1.0 0.5 1.0 (mm)
d
WM
Up
Cell-substrate adhesion
Wound healing
Response to wounding
Ribosomal large subunit biogenesis
Ribosome assembly
Ribonucleoprotein complex biogenesis
Ribosome biogenesis
Cytoplasmic translation
Leukocyte migration
Response to molecule of bacterial origin
ERK1 and ERK2 cascade
Cellular response to lipopolysaccharide
Regulation of inflammatory response
Regulation of hemopoiesis
Response to lipopolysaccharide
Cellular response to lipid
p38MAPK cascade
Down
Synapse pruning
Gliogenesis
Glial cell development
Cell junction disassembly
Glial cell differentiation
Myelination
Ensheathment of neurons
Axon ensheathment
Oligodendrocyte differentiation
Small molecule biosynthetic process
Injury: 3 24 72 (hpi)
p. adjust
Count
e
MG
Regulation of vasculature development
Regulation of angiogenesis
Cell-substrate adhesion
Leukocyte migration
Ribonucleoprotein complex biogenesis
Ribosome assembly
Ribosomal small subunit biogenesis
Ribosome biogenesis
Cytoplasmic translation
Response to wounding
Regulation of neuron death
Regulation of neuron apoptotic process
Positive regulation of neuron death
ERK1 and ERK2 cascade
Regulation of ERK1 and ERK2 cascade
Neurotransmitter transport
Regulation of neurotransmitter levels
Neurotransmitter secretion
Signal release from synapse
Regulation of ion transmembrane transport
Synaptic vesicle cycle
Vesicle-mediated transport in synapse
Injury: 24 72 (hpi)
f
DH
External encapsulating organization
Extracellular structure organization
Extracellular matrix organization
Wounding healing
Ribosome biogenesis
Cytoplasmic translation
Intrinsic apoptotic signaling pathway
Response to wounding
Regulation of apoptotic signaling pathway
Cell chemotaxis
Cellular response to lipid
Regulation of hemopoiesis
Response to molecule of bacterial origin
Response to lipopolysaccharide
Gliogenesis
Glial cell differentiation
Myelination
Ensheathment of neurons
Axon ensheathment
Oligodendrocyte differentiation
Regulation of neurotransmitter levels
Neurotransmitter transport
Neurotransmitter secretion
Signal release from synapse
Vesicle-mediated transport in synapse
Synaptic vesicle cycle
g
VH
Positive regulation of response to external stimulus
Cell-substrate adhesion
Ribosomal large subunit biogenesis
Ribonucleoprotein complex biogenesis
Ribosome assembly
Ribosome biogenesis
Cytoplasmic translation
Response to wounding
Leukocyte migration
Myeloid leukocyte migration
Cellular response to molecule of bacterial origin
p38MAPK cascade
ERK1 and ERK2 cascade
Cellular response to lipopolysaccharide
Regulation of ERK1 and ERK2 cascade
Response to tumor necrosis factor
Import into cell
Sodium ion transport
Import across plasma membrane
Positive regulation of ion transport
Regulation of ion transmembrane transport
Regulation of membrane potential
Neurotransmitter transport
Regulation of neurotransmitter levels

**Extended Data Fig. 3. Analyses and comparisons of different sets of SCI-induced DEGs**

(**a-c**) The bar graphs showing the up- and down-regulated DEGs at each anatomic region (**a**), directionality (**b**) or distance to the injury site (**c**) at the indicated time points after SCI, compared to the group of 0 hpi. Note that the WM and the MG show more anatomic domain-specific DEGs than the other two regions, while directionality or distance to the injury site exhibits minimal differential effects except for the 0.5 mm sections at 72 hpi.

(**d-g**) The top 5 GO terms of the up- or down-regulated DEGs associated with the WM (**d**), MG (**e**), DH (**f**) and VH (**g**) of the spinal cord are shown as dot plots at the indicated time points after SCI. GO terms involving early injury responses such as the ERK1/2 signaling cascade and immune and inflammation activation such as leukocyte migration and response to lipopolysaccharide (LPS) are the most frequently observed terms within 72 hpi across all four anatomic regions, reflecting the major molecular events in the immediate and the acute phases of SCI.

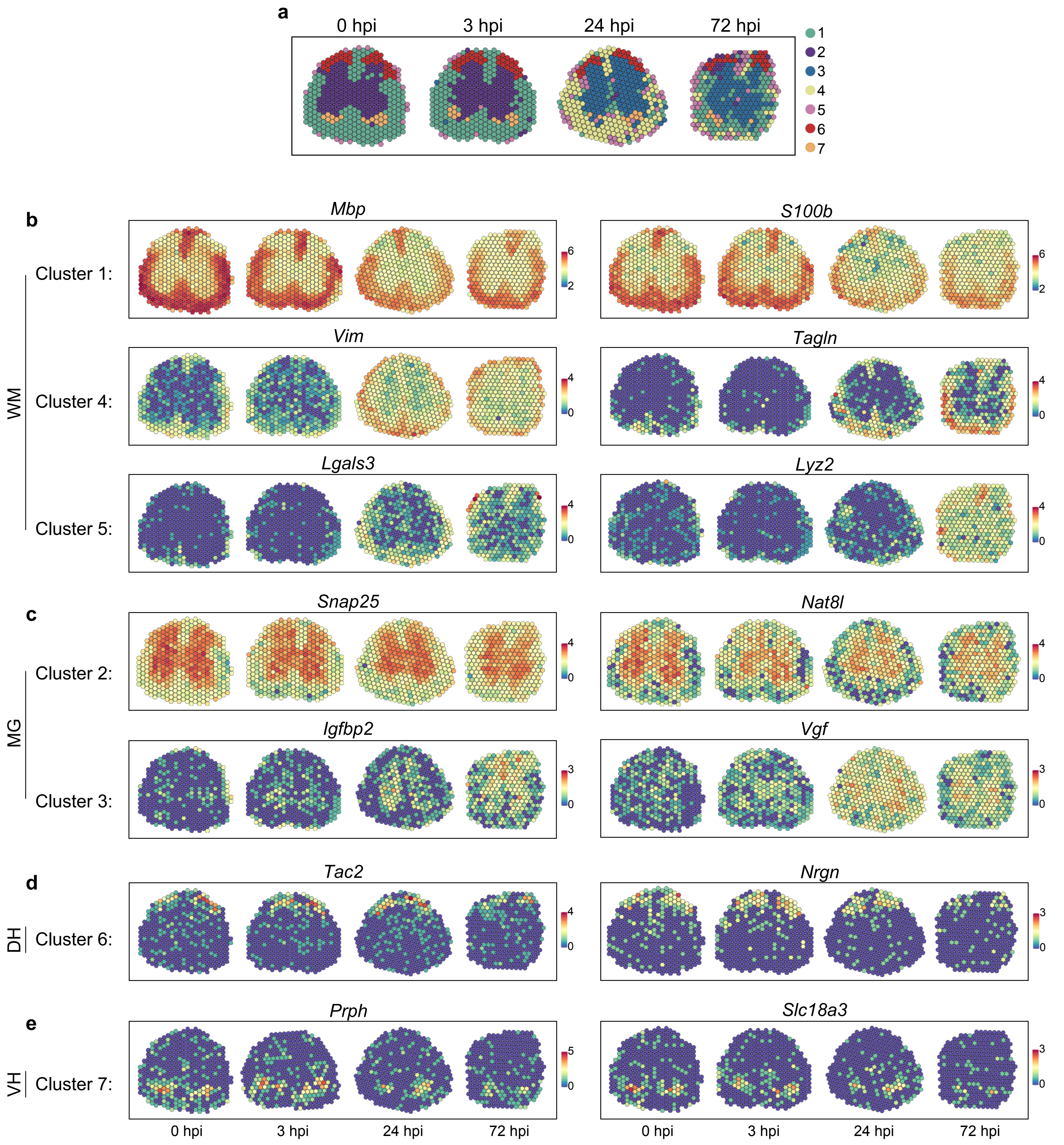
a
0 hpi
3 hpi
24 hpi
72 hpi
1
2
3
4
5
6
7
b
WM
Cluster 1:
Cluster 4:
Cluster 5:
Mbp
S100b
Vim
Tagln
Lgals3
Lyz2
c
MG
Cluster 2:
Cluster 3:
Snap25
Nat8l
Igfbp2
Vgf
d
DH
Cluster 6:
Tac2
Nrgn
e
VH
Cluster 7:
Prph
Slc18a3
0 hpi
3 hpi
24 hpi
72 hpi
0 hpi
3 hpi
24 hpi
72 hpi

**Extended Data Fig. 4. The spatial views of the representative genes of the 7 spatiotemporal clusters**

(**a**) The spatial distribution of the 7 spatiotemporal clusters in the mouse spinal cord (-1.0 mm section) at the indicated time points of SCI.

(**b-e**) The spatial views of the expression level of the representative marker genes of Cluster 1 (*Mbp*, *S100b*), Cluster 4 (*Vim*, *Tagln*) and Cluster 5 (*Lgals3*, *Lyz2*) in the WM (**b**), Cluster 2 (*Snap25*, *Nat8l*) and Cluster 3 (*Igfbp2*, *Vgf*) in the MG (**c**), Cluster 6 (*Tac2*, *Nrgn*) in the DH (**d**), and Cluster 7 (*Prph*, *Slc18a3*) in the VH (**e**) of the mouse spinal cord (-1.0 mm section) at the indicated time points of SCI.

Extended Data Fig. 5 - SCI ST

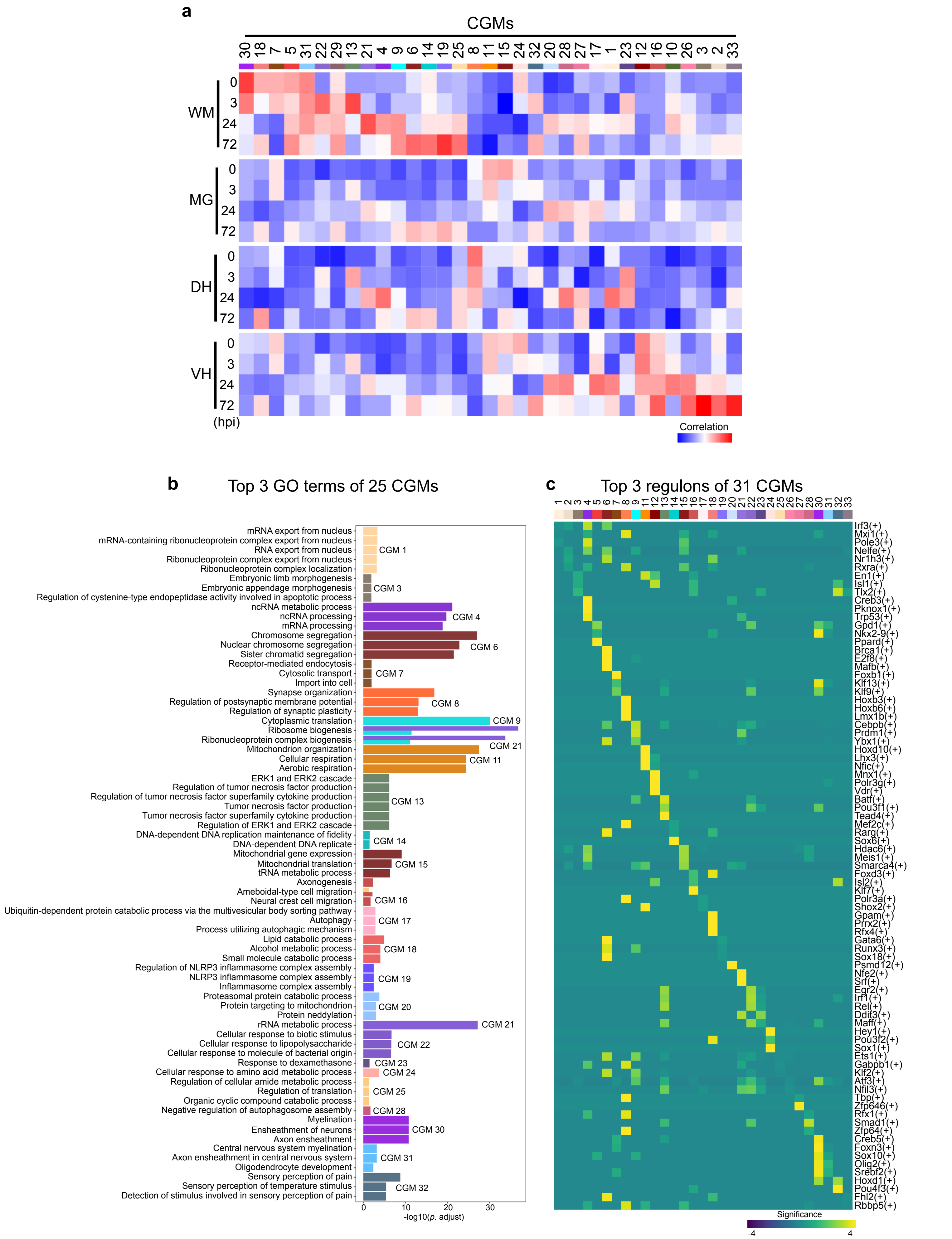

**Extended Data Fig. 5. The overview of the 33 co-expressed gene modules (CGMs) in SCI**

(**a**) The Pearson correlation coefficients between each CGM and the four anatomic domains at the indicated time points of SCI.

(**b**) The top 3 ($p < 0.05$) enriched GO terms (including tied ranks) of the 25 CGMs. The other 8 CGMs are not enriched significantly with any specific GO term and are thus not shown.

(**c**) The top 3 regulons (ordered by significance derived from the hypergeometric test, see Methods) of 31 CGMs. The other two CGMs are not enriched significantly with any specific regulon and are thus not shown.

### Extended Data Fig. 6a- SCI ST

**a**

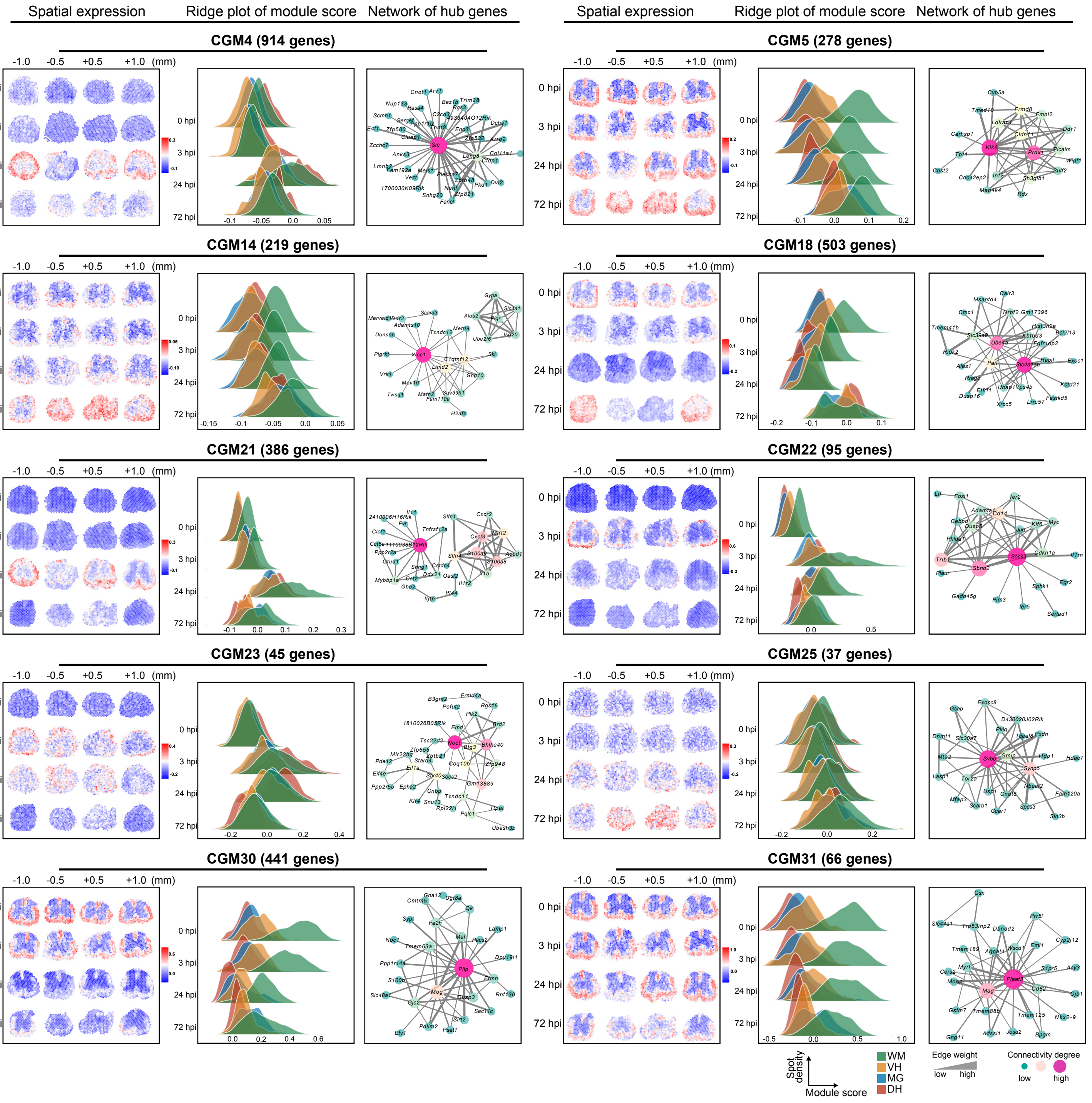

### Extended Data Fig. 6b - SCI ST

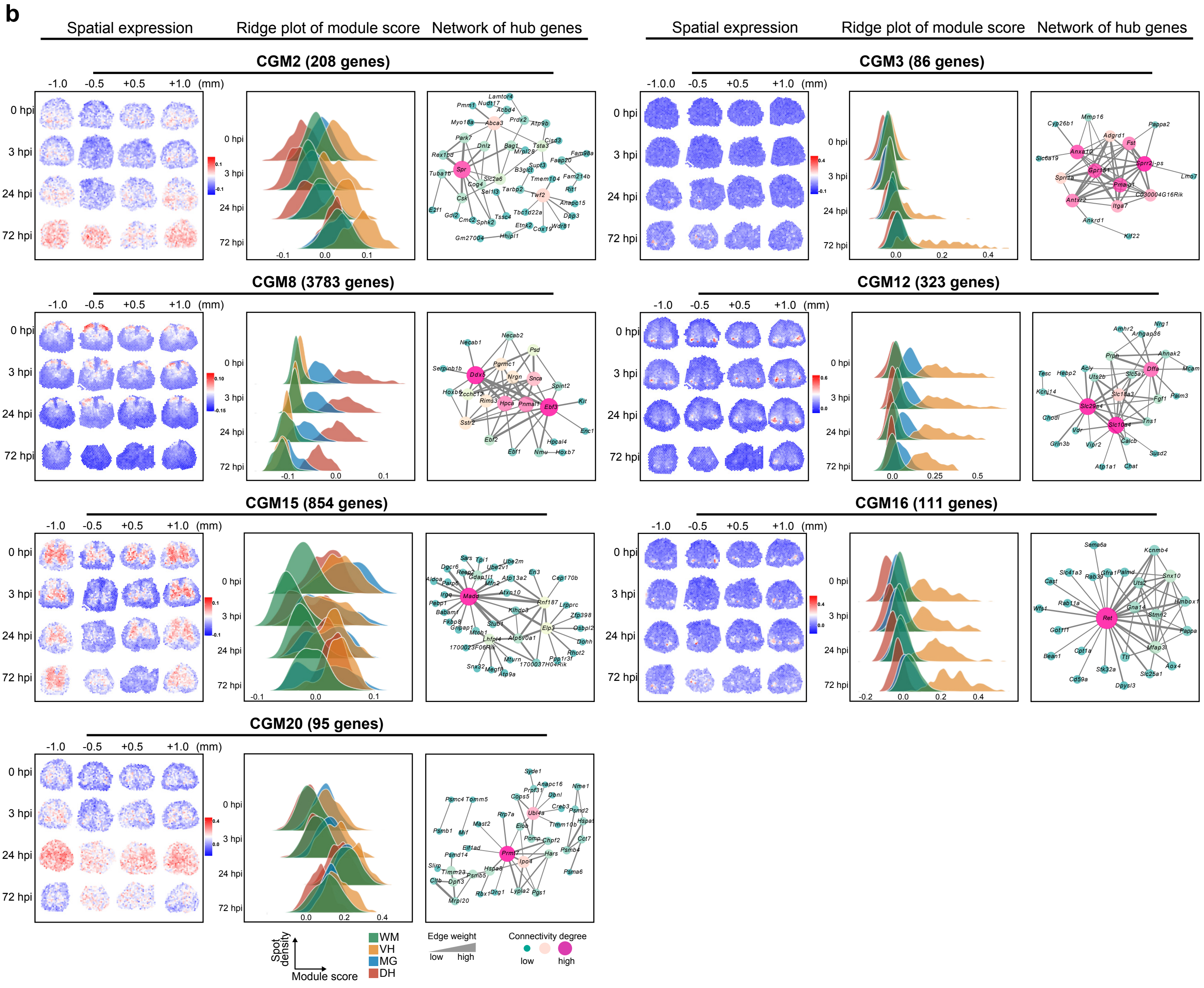

### Extended Data Fig. 6c - SCI ST

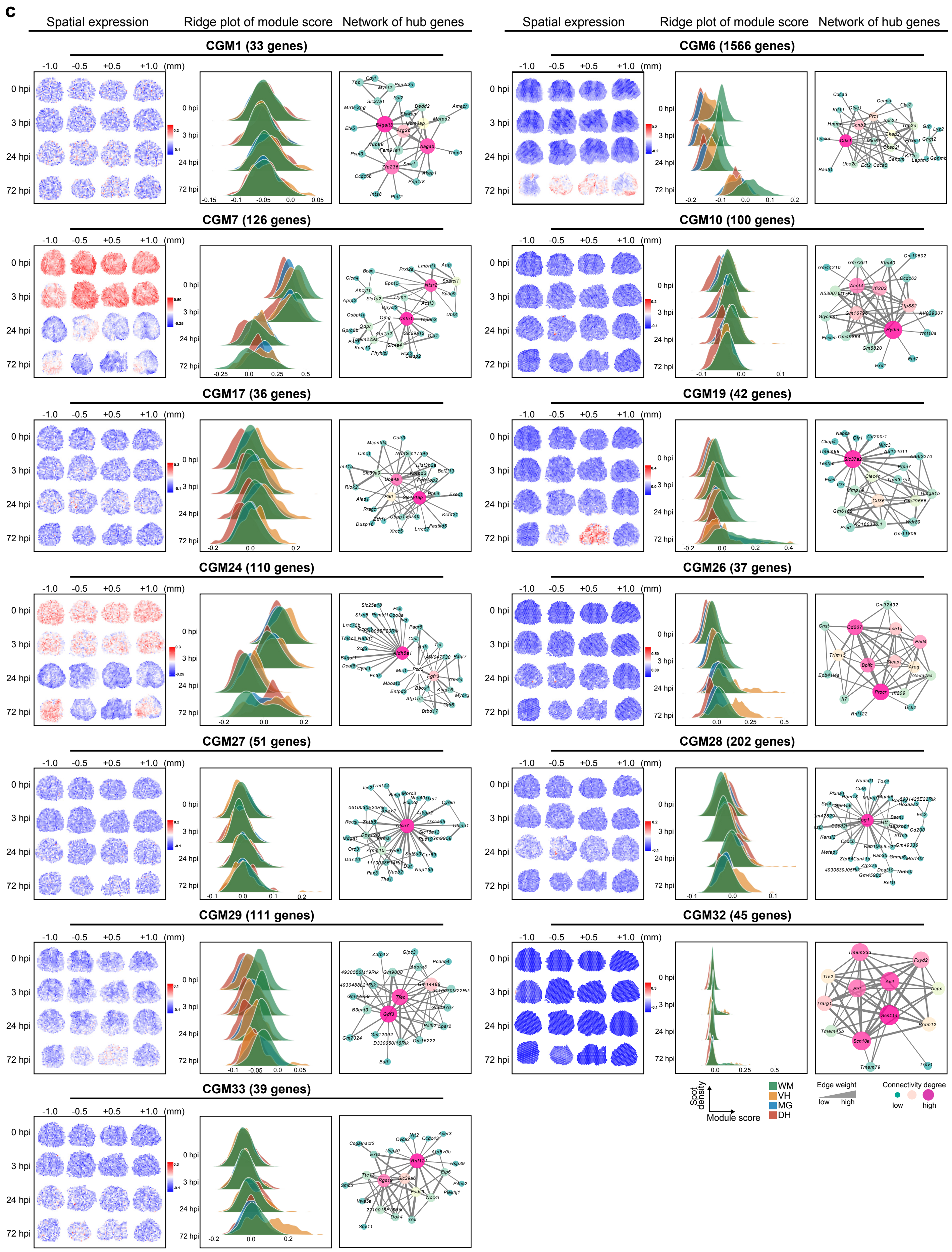

**Extended Data Fig. 6. The spatiotemporal dynamics and regulatory networks of the CGMs**

Analysis of the molecular features of the rest 30 CGMs. From left to right: left, the spatial view of average levels of the genes expressed in the CGM at the indicated time points of SCI; middle, the ridge plots quantitatively illustrating the density distribution of the module gene expression within the 4 anatomic domains; right, the representative GO terms (top) and the hub gene network showing the highest 50 connectivity weights (bottom) of the indicated CGM. The number of genes in each CGM is shown in the parentheses.

(**a**) Ten WM-associated CGMs, displaying the spatial changes in gene expression predominantly in the WM. For example, CGM22 shows an increased expression in the WM specifically at 3 hpi, which is similar to that of CGM13 in Fig. 2c but of the different hub genes *Socs3* and *Sbno2*, and the GO term analysis (see Extended Data Fig. 6a) suggests CGM13 plays an anti-inflammatory role, likely by inhibition of the phosphorylation of *Stat3*; CGM30 and CGM31 are related to axonal myelination in the WM and their gene expressions are substantially decreased at 24 and 72 hpi, respectively, suggesting the onset of demyelination at 24 hpi and/or afterwards; CGM21 shows an increase of gene expression specifically in the WM at 24 hpi, which matches the spatiotemporal pattern of the Cluster 4 (Fig. 1e, k-l) and may be involved in regulating inflammatory responses especially by stimulating leukocyte recruitment; CGM25 is initially increased in the WM at 24 hpi and then also in the GM at 72 hpi (especially at the ±0.5 mm sections), and the hub genes of CGM25 contain *Svbp*, which encodes a tubulin carboxypeptidase that regulates axon and synapse formation as well as neuron differentiation, implying an induction or an attempt of induction of neuro-regenerative programs at the transition of acute to subacute phase of SCI.

(**b**) Seven GM-associated CGMs. For example, CGM8 (DH), CGM12 (VH) and CGM16 (VH) exhibit DH- or VH-specific expression patterns and are related to neuronal functions such as synapse organization and ion transport; CGM20 and CGM2 both show increased gene expression in MG at 24 and 72 hpi, respectively, and CGM20 is functionally associated with vesicle transport whereas CGM2 is mostly involved in the proteasome protein catabolic process.

(**c**) Thirteen CGMs without obvious domain-enriched expression changes. For example, CGM24 associated with organic acid catabolic process is highly expressed at all spinal cord sections at 0 and 3 hpi, but drastically decreased afterwards, likely reflecting and/or due to demyelination; CGM6 related to chromosome segregation is remarkably increased in the spots spanning both the WM and the GM at 72 hpi, suggesting widespread, active DNA replication and cell proliferation by the end of the acute phase of SCI.

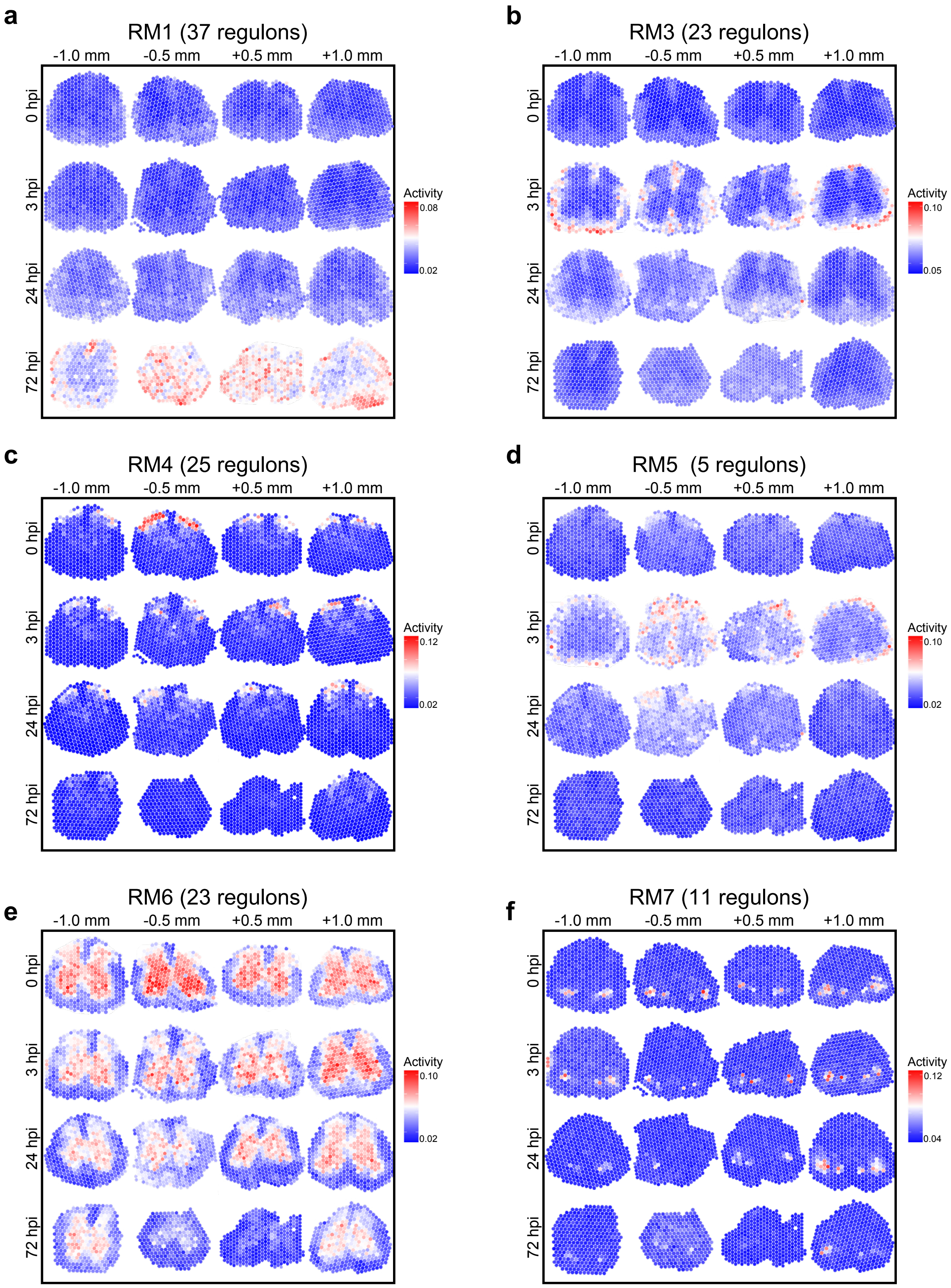
a
RM1 (37 regulons)
-1.0 mm
-0.5 mm
+0.5 mm
+1.0 mm
0 hpi
3 hpi
24 hpi
72 hpi
Activity
0.08
0.02
b
RM3 (23 regulons)
Activity
0.10
0.05
c
RM4 (25 regulons)
Activity
0.12
0.02
d
RM5 (5 regulons)
Activity
0.10
0.02
e
RM6 (23 regulons)
Activity
0.10
0.02
f
RM7 (11 regulons)
Activity
0.12
0.04

**Extended Data Fig. 7. The spatiotemporal activity of the regulon modules (RMs) in SCI**

(**a-f**) The spatial views of the activity of the rest 6 RMs in SCI. The number of the transcriptional regulons in each RM is shown in the parentheses. The RMs are highly correlated with the anatomic domain and injury time. For example, RM1 is activated specifically at 72 hpi (**a**); RM3 (**b**) and RM5 (**d**) are highly active in the WM at 3 hpi; RM4 (**c**) and RM7 (**f**) are DH- and VH-associated, respectively; and RM6 (**e**) is predominantly associated with the MG and its activity decreased along the time after SCI.

**Extended Data Fig. 8- SCI ST**

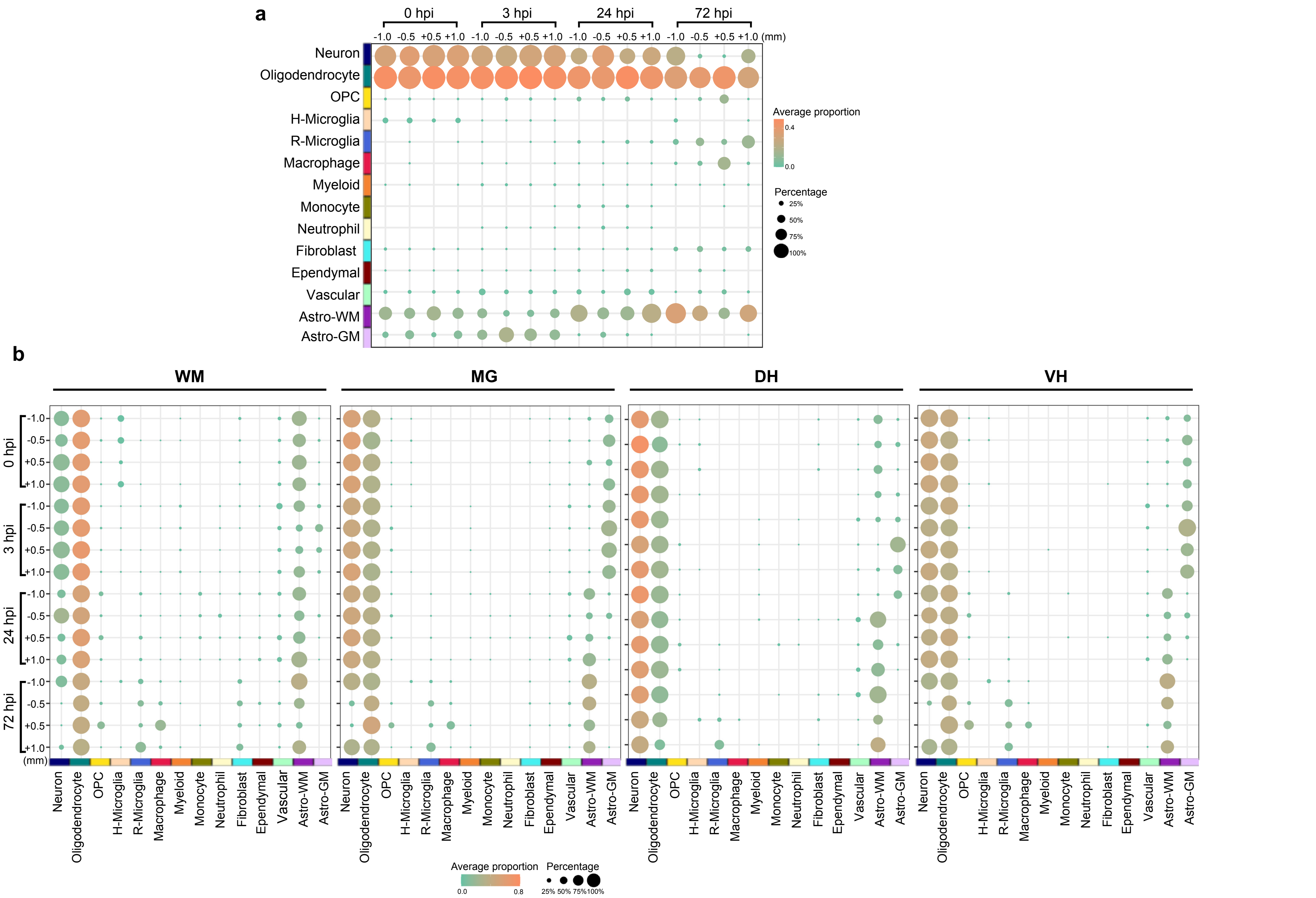

**Extended Data Fig. 8. The quantifications of the spinal cell type deconvolution in relation to the anatomic domain and injury time**

The dot plots showing the average proportion of the indicated cell type in each spatial spot (color of the dot) and the percentage of the spatial spots containing the indicated cell type (size of the dot) throughout the entire spinal cord section (**a**) or in the specific anatomic domains (**b**) at indicated time points of SCI.

#### Extended Data Fig. 9 - SCI ST

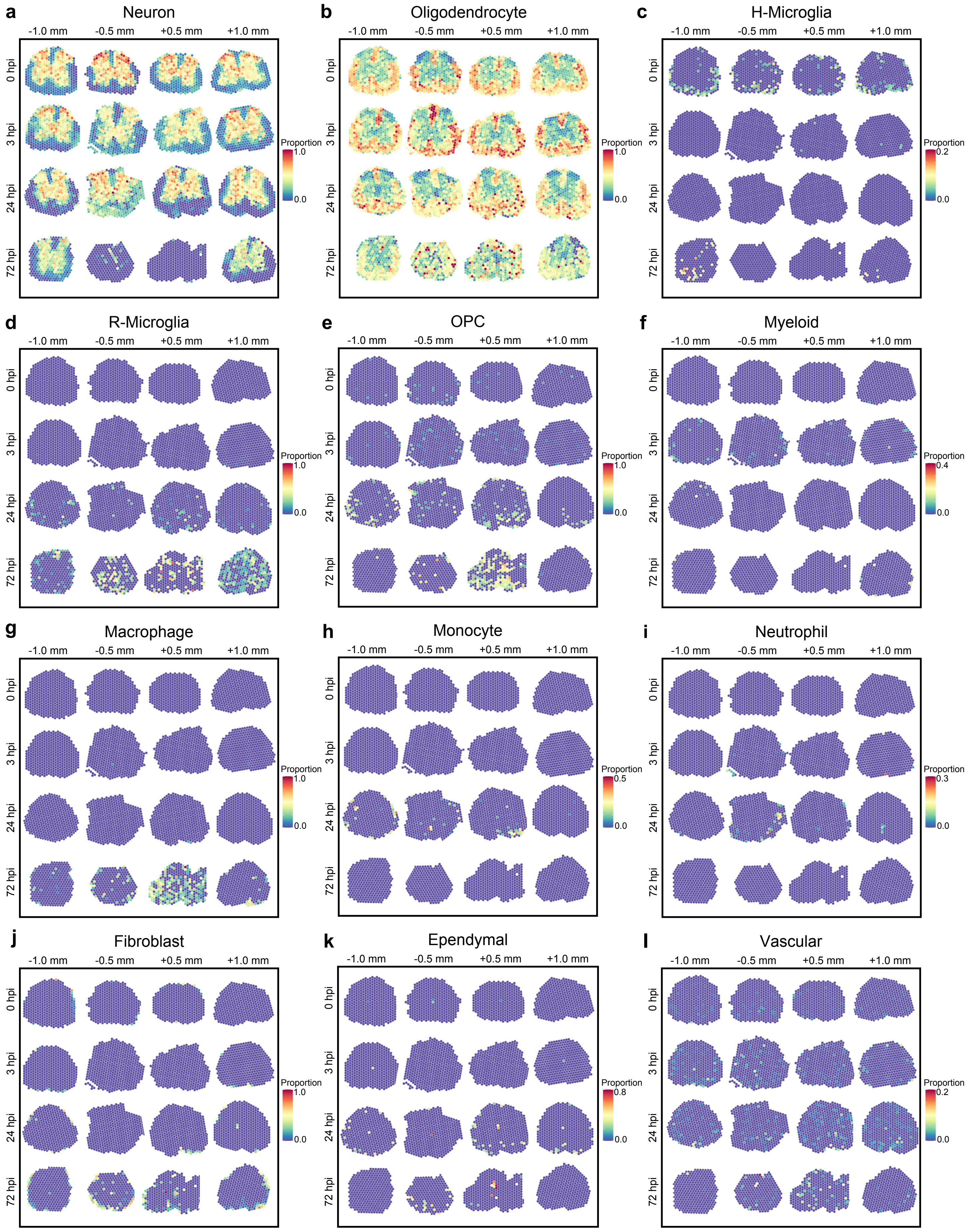

**Extended Data Fig. 9. The spatiotemporal distributions of spinal cell types in SCI**

(**a, b**) Both neurons in the GM (**a**) and oligodendrocytes in the WM (**b**) decreased slightly at 24 hpi and drastically at 72 hpi, indicative of injury-induced neurodegeneration and demyelination.

(**c, d**) Homeostatic (H)-microglia were substantially reduced at 3 hpi (**c**), whereas reactive (R)-microglia emerged at 24 and 72 hpi (**d**).

(**e**) Oligodendrocyte precursor cells (OPCs) were increased along the time after SCI, which might represent a compensatory myelinogenesis.

(**f-i**) The spatiotemporal distributions of several immune and inflammation-related cell types, including myeloid cells (**f**), macrophages (**g**), monocytes (**h**) and neutrophils (**i**).

(**j-l**) Fibroblasts (**j**), ependymal cells (**k**) and vascular cells (**l**) previously shown to participate in the scar formation were activated in SCI. Noteworthily, our analysis allocated ependymal cells only in one or two spatial spots at the very center of the intact spinal cord, which faithfully reflected the anatomic localization and function of ependymal cells in lining the central canal of the spinal cord, and developed a diverse astroependymal distribution later in SCI.

### Extended Data Fig. 10 - SCI ST

**a**

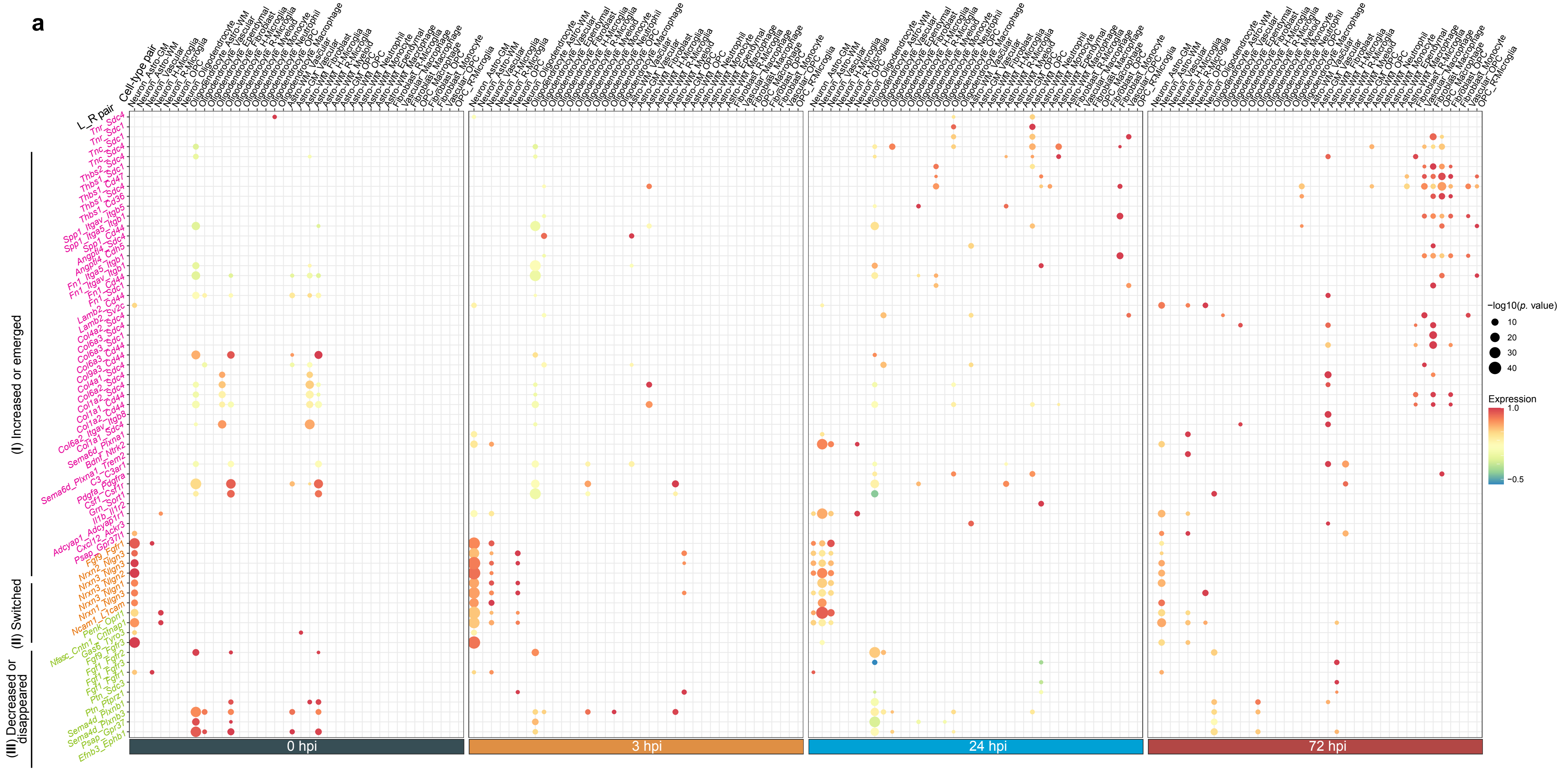


**b**

| Expression of the L_R pairs | Biological functions |
|---|---|
| Increased or emerged upon injury (I) | Extracellular matrix organization and fibrosis<br>Tube formantion and epithelial-mesenchymal transformation<br>Inflammation and immune response<br>Chemotaxis of macrophage and macrophage activition |
| Switched from one cell-type pair to another (II) | Neural cell adhesion<br>Synapse adhesion and dendrotogenesis<br>Axon guidance and actin remodeling<br>Neurotrophic and neuroprotective |
| Decreased or disappeared upon injury (III) | Axon growth and guidance<br>Dendritic spine development<br>Synapse formation/elimination<br>Myelination |

**c**

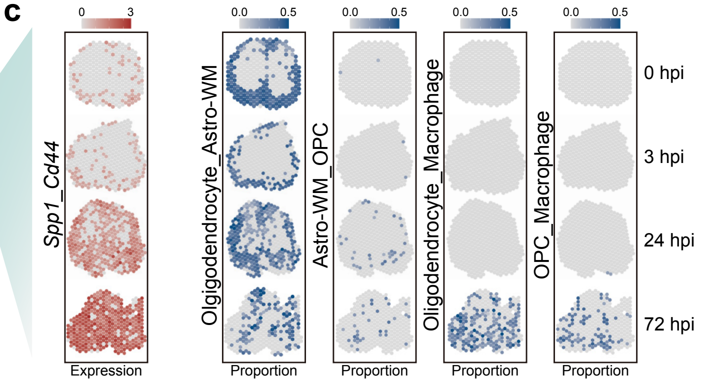

**Extended Data Fig. 10. The SA-CCC analysis of *in situ* multicellular communications and the L_R pairs mediating the cell-cell interactions in SCI**

(**a**) Dot plots showing the top 2 (*p*. value) interacting L_R pairs in each cell-type pair at the indicated time points in SCI. Color denotes the average expression levels of each L_R pair in the indicated, colocalized cell-type pairs.

(**b**) According to their changes in SCI, the L_R pairs in (a) are categorized into three main classes: (Class-I) the expression of the L_R pairs increases or emerges upon injury (e.g., the *Col_Sdc* families), which mainly participate in cell matrix adhesion, cell migration, immune, angiogenesis and wound healing; (Class-II) the expression of the L_R pairs switches from one cell-type pair to another (e.g., the *Nrxn_Nlgn* families), which are involved in neuron and synapse adhesion, dendritogenesis, actin remodeling, and neuroprotection; and (Class-III) the expression of the L_R pairs decreases or disappears (e.g., the *Fgf_Fgfr* signaling pathway), which regulate axon growth and guidance, dendritic spine development, synapse formation and elimination, and myelination

(**c**) Left: the spatial views of the interaction of a representative Class-I L_R pair *Spp1_Cd44* in (a); right: the spatial distributions of the major cell-type pairs hosting the L_R pair of *Spp1_Cd44* along the time during SCI.

### Extended Data Fig. 11 - SCI ST

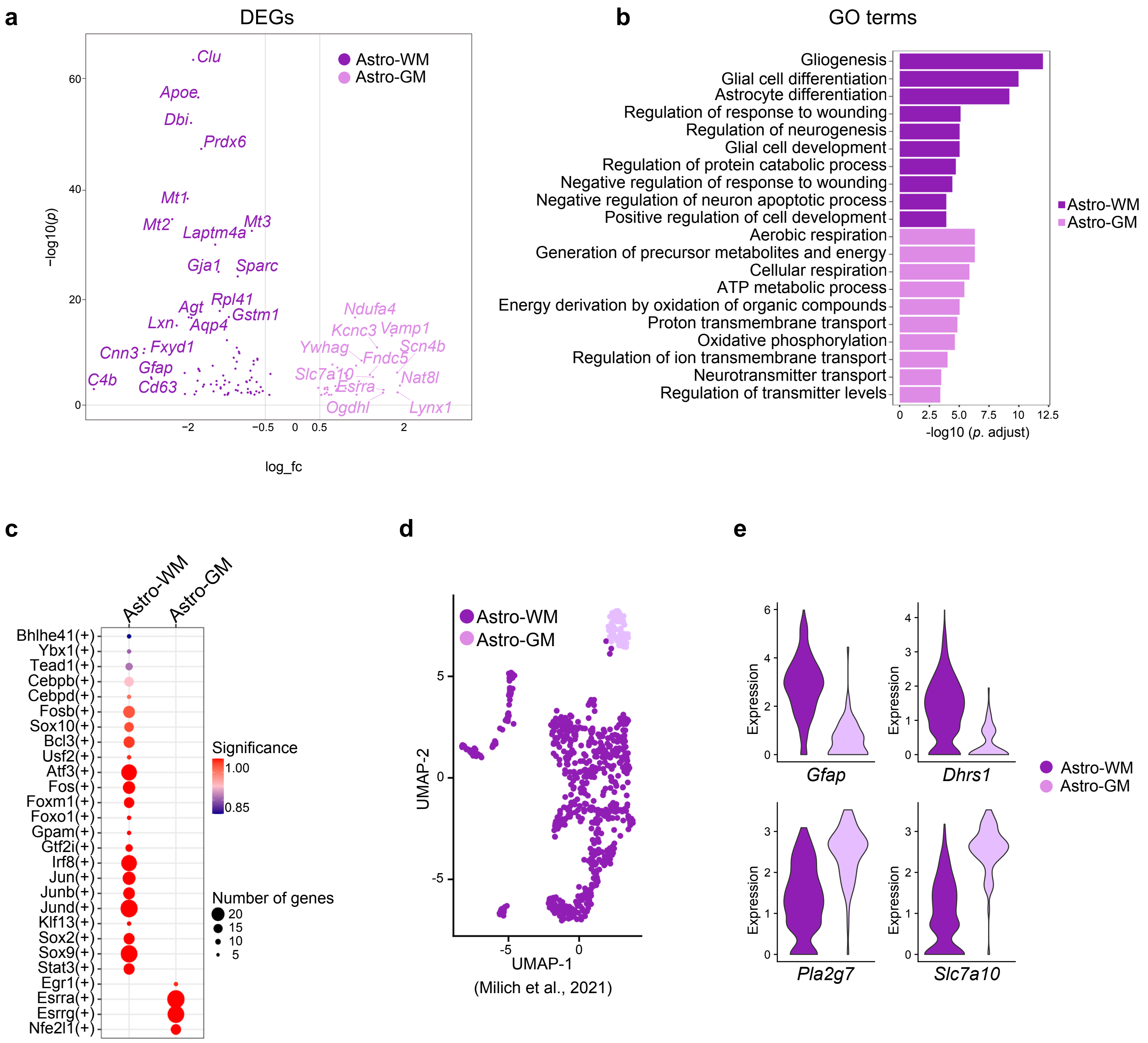

**Extended Data Fig. 11. Comparisons of the molecularly-defined astrocytes-WM and -GM in the mouse spinal cord**

(**a-c**) The volcano plots showing the DEGs, analyzed by cell type-specific inference of differential expression (C-SIDE) (**a**), the top 10 (*p*. value < 0.05) GO terms (**b**), and the enriched regulons (**c**) of the DEGs of astrocytes-WM and -GM of the mouse spinal cord at 0 hpi. Astrocytes-WM were highly associated with the function of gliogenesis, glial cell development and differentiation, inhibition of neuronal apoptosis and wound response, with the TFs of the main regulons (e.g., Jund, Sox9, Atf3 and Irf8) enriched in cell proliferation, differentiation and death, and cellular stress and immune responses[81-83]; astrocytes-GM were energetically active and provided trophic supports to neurons by mediating transmembrane transport of ions, metabolites and neurotransmitters, with the TFs of the main regulons (e.g., Esrra and Esrrg) enriched in cellular energy production[84,85].

(**d**) The re-analysis of the astrocytes in the SCI single-cell dataset[9] by UMAP reveals distinct astrocyte-WM and -GM clusters.

(**e**) The violin plots showing the expression levels of the representative marker genes of *Gfap* and *Dhrs1* (for astrocytes-WM) and *Pla2g7* and *Slc7a10* (for astrocytes-GM).

#### Extended Data Fig. 12 - SCI ST

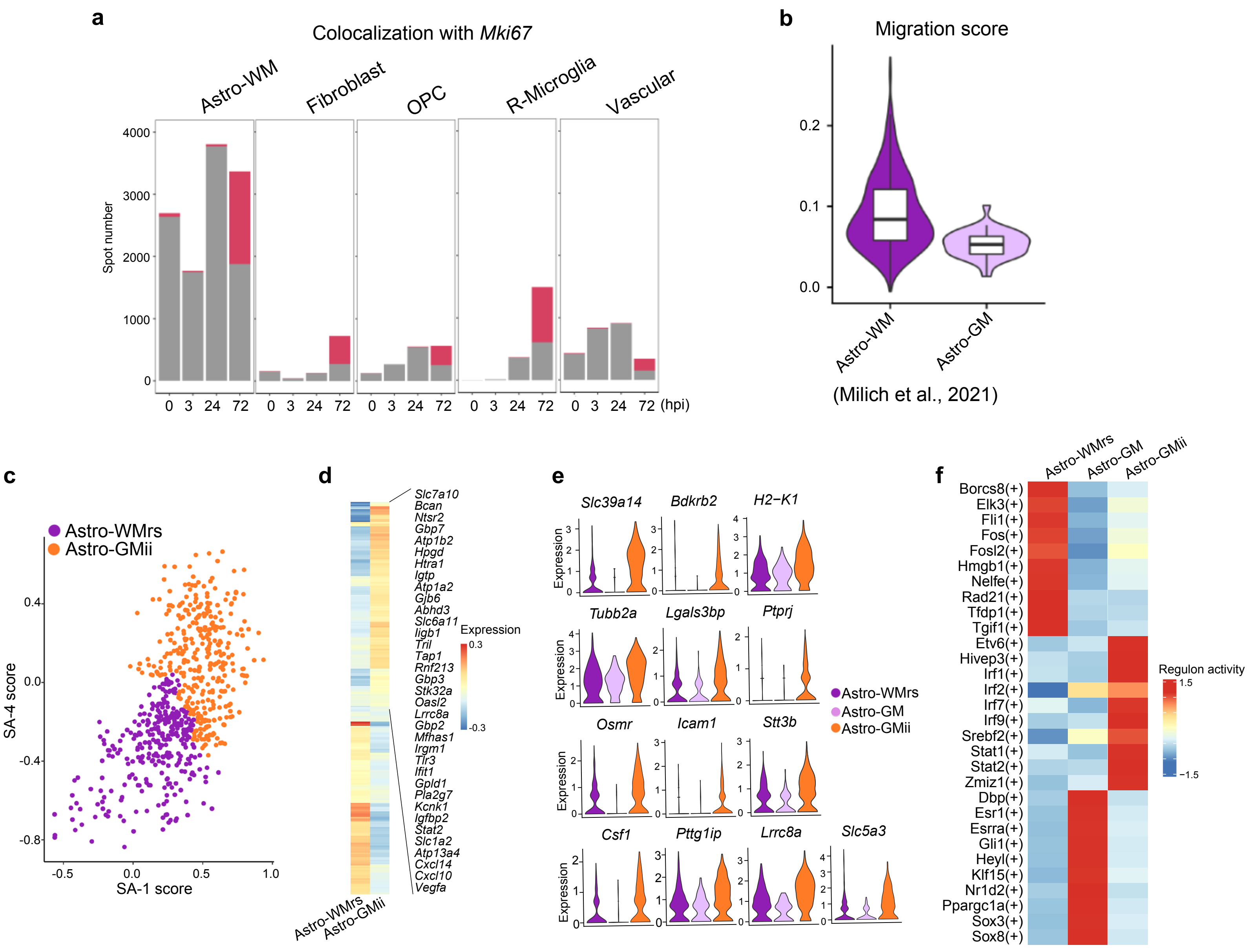

**Extended Data Fig. 12. Characterization of the molecular features of the spatiotemporally-defined astrocytes-GMii in SCI**

(**a**) Quantifications of the numbers of the spatial spots located in the high proliferation score regions (colocalization with *Mki67* expression) (red) and the total spatial spots containing the corresponding cell types (grey and red) at the indicated time points of SCI.

(**b**) The violin plot showing the migration score of the astrocytes-WM and -GM in the SCI single-cell dataset[9].

(**c**) Astrocytes-*Gfap* are classified into two domain-associated states using the scores of the SA-1 and SA-4 gene sets identified in Fig. 4g.

(**d**) The heatmap of the marker genes of astrocytes-WMrs and -GMii identified in (c) and the representative upregulated genes denoting astrocytes-GMii are shown.

(**e**) The violin plots showing the potential surface marker genes that are predominantly expressed in astrocytes-GMii compared to astrocytes-WMrs and -GM.

(**f**) The heatmap showing the top 10 ($p < 0.05$) enriched regulons of astrocytes-GMii, -GM and -WMrs based on the analysis of the SCI single-cell dataset.

Astro, astrocytes; GM, grey matter; WM, white matter; Astro-GMii, injury-induced GM-relocated astrocytes; Astro-WMrs: the rest astrocytes-WM (which together with Astro-GMii constitute the astrocyte-*Gfap* population).

**Extended Data Fig. 13 - SCI ST**

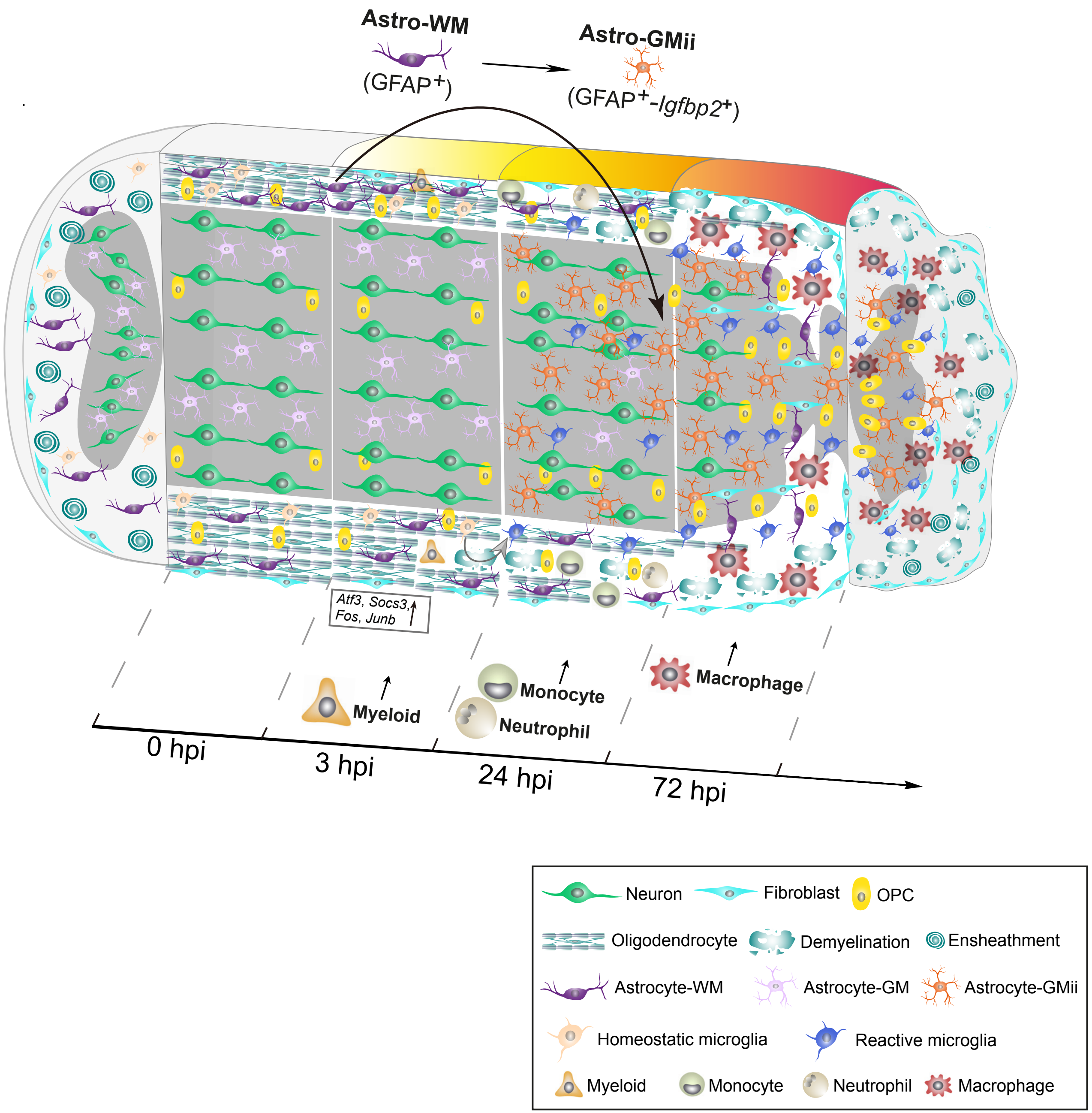

**Extended Data Fig. 13. A schematic diagram of SCI-induced spatiotemporal landscape**

Upon SCI, a cluster of early response genes are activated at 3 hpi, accompanied by induction of immune and inflammatory responses and elimination of homeostatic microglia in the WM, which are followed by demyelination in the WM and loss of neurons and astrocytes in the GM starting from 24 hpi. Meanwhile, a variety of immune cells are activated and recruited to the injury site. For example, myeloid cells are induced at 3 hpi; monocytes and neutrophils infiltrate into the WM at 24 hpi; reactive microglia increase dramatically at 24 and 72 hpi; and macrophages are found at 72 hpi. Fibroblasts are usually located at the periphery of the intact spinal cord but show up in the GM at 72 hpi in SCI. Astrocytes in the WM are also activated, of which a population of *Igfbp2*$^+$-GFAP$^+$ astrocytes migrate into the GM and become astrocytes-GMii. These cells interact with neurons in the GM at 24 and 72 hpi and play roles in supporting neuro-/synaptogenesis and regulating sterol and steroid biosynthesis. Thus, they likely functionally compensate for the loss of the original astrocytes-GM. Note that the gradient of color from white to orange and red indicates increasing complexity of SCI-induced cell changes and severity of tissue damage along the time after injury (from left to right).

#### SUPPLEMENTAL TABLES

**Supplemental Table 1. The mouse sample information for the SCI spatial RNA-seq.**

**Supplemental Table 2. The genes included in each CGM.**

**Supplemental Table 3. The regulons included in each RM.**

**Supplemental Table 4. The gene sets for the annotation of astrocytes-WM and GM.**

**Supplemental Table 5. The gene sets of the four SA groups.**

**Supplemental Table 6. The unique and shared DEGs of astrocytes-WM, -GM and -GMii.**
